# mRNA-1273 efficacy in a severe COVID-19 model: attenuated activation of pulmonary immune cells after challenge

**DOI:** 10.1101/2021.01.25.428136

**Authors:** Michelle Meyer, Yuan Wang, Darin Edwards, Gregory R. Smith, Aliza B. Rubenstein, Palaniappan Ramanathan, Chad E. Mire, Colette Pietzsch, Xi Chen, Yongchao Ge, Wan Sze Cheng, Carole Henry, Angela Woods, LingZhi Ma, Guillaume B. E. Stewart-Jones, Kevin W. Bock, Mahnaz Minai, Bianca M. Nagata, Sivakumar Periasamy, Pei-Yong Shi, Barney S. Graham, Ian N. Moore, Irene Ramos, Olga G. Troyanskaya, Elena Zaslavsky, Andrea Carfi, Stuart C. Sealfon, Alexander Bukreyev

**Author notes:** Address correspondence to: Andrea Carfi, Moderna Inc, Cambridge, MA 02139, Stuart C. Sealfon, Icahn School of Medicine at Mount Sinai, New York, NY 10029, USA. Alexander Bukreyev, Departments of Pathology and Microbiology & Immunology, Galveston National Laboratory, University of Texas Medical Branch, Galveston, Texas 77555-0609, USA. These authors contributed equally (Michelle Meyer, Yuan Wang, Darin Edwards, Gregory R. Smith, Aliza B. Rubenstein, Palaniappan Ramanathan).

## Abstract

The mRNA-1273 vaccine was recently determined to be effective against severe acute respiratory syndrome coronavirus 2 (SARS-CoV-2) from interim Phase 3 results. Human studies, however, cannot provide the controlled response to infection and complex immunological insight that are only possible with preclinical studies. Hamsters are the only model that reliably exhibit more severe SARS-CoV-2 disease similar to hospitalized patients, making them pertinent for vaccine evaluation. We demonstrate that prime or prime-boost administration of mRNA-1273 in hamsters elicited robust neutralizing antibodies, ameliorated weight loss, suppressed SARS-CoV-2 replication in the airways, and better protected against disease at the highest prime-boost dose. Unlike in mice and non-human primates, mRNA-1273- mediated immunity was non-sterilizing and coincided with an anamnestic response. Single-cell RNA sequencing of lung tissue permitted high resolution analysis which is not possible in vaccinated humans. mRNA-1273 prevented inflammatory cell infiltration and the reduction of lymphocyte proportions, but enabled antiviral responses conducive to lung homeostasis. Surprisingly, infection triggered transcriptome programs in some types of immune cells from vaccinated hamsters that were shared, albeit attenuated, with mock-vaccinated hamsters. Our results support the use of mRNA-1273 in a two-dose schedule and provides insight into the potential responses within the lungs of vaccinated humans who are exposed to SARS-CoV-2.

## INTRODUCTION

When the World Health Organization declared SARS-CoV-2 a global pandemic in March 2020, Phase 1 clinical trials on the most promising vaccine candidates had already commenced. Nucleoside modified mRNA, a relatively new addition to the arsenal of vaccine platforms, has shown promise against numerous viral infectious diseases in preclinical trials (1, 2), and phase 1 and 2 trials that are completed (3, 4) or currently underway (5–7). Previous preclinical work on a related betacoronavirus enabled the rapid development of mRNA-1273, a vaccine composed of a modified mRNA encoding for a stabilized prefusion form of the SARS-CoV-2 spike (S) protein encapsulated in lipid nanoparticles (1). Vaccination with mRNA-1273 prevented infection in the lungs and upper airways of mice (1) and rhesus macaques (8). In phase 1 (9, 10) and phase 2 studies the vaccine was found to be safe and to elicit robust neutralizing antibody responses. More recently, interim analysis of Phase 3 data indicated mRNA-1273 was 94.1% efficacious in prevention of COVID-19 disease and highly efficacious in protecting against severe disease (11). On December 18, 2020, the U.S. Food and Drug Administration authorized the emergency use of mRNA-1273.

Nonhuman primate (12–14), ferret (15–17) and mouse (18) models that have been developed to understand SARS-CoV-2 pathogenesis and test vaccine and therapeutics efficacies display a mild course of disease. The lack of overt symptoms in NHPs and ethical constraints due to the sheer numbers required to recapitulate low frequency human cases with severe clinical manifestations limits the ability to understand all aspects of the disease and vaccine effectiveness. Hamsters are highly susceptible to SARS-CoV-2 and develop a severe pneumonia similar to COVID-19 patients (19–22). Hamster angiotensin converting enzyme – 2 (ACE2) receptors, that enable SARS-CoV-2 binding and entry, are analogous to the human receptor(20) and therefore, virus adaptation (18) or transgenic modifications (23) are not required for permissiveness. Additionally, hamsters can support high levels of virus replication and transmission, and develop severe clinical symptoms, weight loss and lethality with high SARS-CoV-2 infectious doses (21), making them an ideal model for evaluating vaccines.

Here, we tested the efficacy of the mRNA-1273 vaccine with prime only and three dose levels of prime-boost regimens using the stringent golden Syrian hamster model. Serological, virological, clinico-pathological and single cell (sc) RNA-seq analyses were conducted to characterize vaccine-mediated immunity before and after challenge. We show that vaccination induced robust virus neutralizing antibody responses, attenuated virus replication, and mitigated against the influx of inflammatory innate immune cells and the relative reduction of lymphocytes in the lungs after challenge, although activated immune cell populations were observed. Our data illustrates the viral, cellular, and immune dynamics within the lungs of vaccinated hamsters which may offer a unique perspective into the events which occur within the lungs of vaccinated humans who are exposed to SARS-CoV-2.

## RESULTS

### Vaccination induces potent antibody responses

Three groups of outbred hamsters (*n* = 15) were vaccinated with 25 µg, 5 µg and 1 µg of mRNA-1273 via the intramuscular (IM) route in a prime (week 0)-boost (week 3) regimen (Figure 1A). One group of hamsters (*n* =15) received a prime-only dose of 25 µg at week 0 and another group (*n* =15) were mock vaccinated as the study’s control. The humoral responses to vaccination were measured by enzyme-linked immunosorbent assay (ELISA) specific for the SARS-CoV-2 S protein (Figure 1B) and its receptor binding domain (RBD; Figure 1C). Three weeks after the prime dose, higher S-specific IgG titers were detected in hamsters vaccinated with 25 µg and 5 µg doses compared to the 1 µg dose (Figure 1B). S-specific IgG titers were significantly augmented in all groups following the booster, but continued to be significantly higher in the 25 µg and 5 µg dose groups compared to 1 µg dose group. RBD-specific IgG titers were comparable among hamsters in the prime-boost regimen groups after receipt of their first dose (Figure 1C). After a boost dose, RBD titers had increased significantly (*P*<0.0001) and remained comparable among the prime-boost groups. RBD titers in the prime-boost vaccine groups were also significantly higher after the second dose when compared to the 25 µg prime-only group (*P*<0.0001), while titers remained unchanged for the 25 µg prime-only group between weeks 3 and 6 post-vaccination. The ability of serum to neutralize live SARS-CoV-2 reporter virus was also determined (Figure 1D). Most vaccinated hamsters produced neutralizing titers after the prime dose. As with RBD-binding titers, neutralizing titers significantly increased in all groups which received a booster dose, and the levels remained comparable among these hamster groups, but higher versus the prime-only vaccine group. The magnitude of neutralizing antibody titers in all booster-vaccinated groups was higher than in convalescent COVID-19 patients, while titers in the prime-only group were comparable to those seen in these subjects. Neutralizing titers significantly correlated with S-specific IgG titers (Supplemental Figure 1A) and RBD-specific titers (Supplemental Figure 1B), albeit the correlation was greater with RBD-specific IgG titers at both weeks 3 and 6 post-vaccination

**Figure 1.**
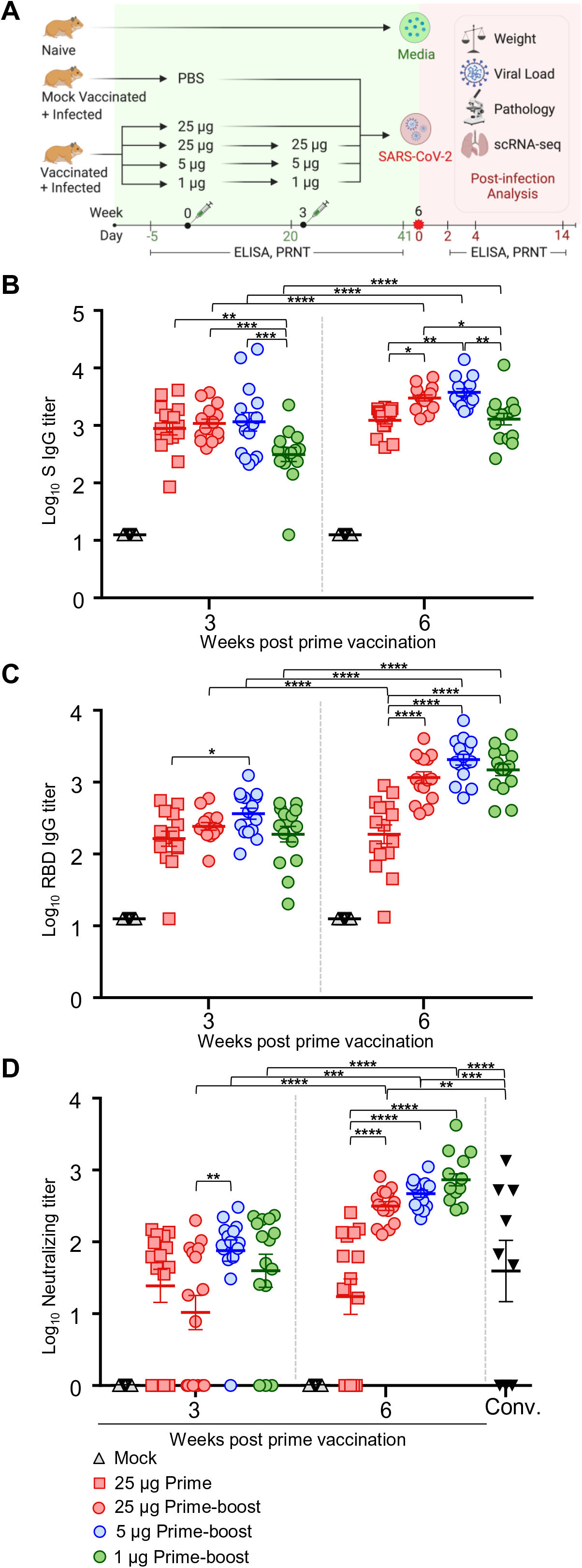
Serum antibody responses in vaccinated hamsters. (A) Study design. Hamsters were prime vaccinated via the IM route at week 0 and boosted at week 3, with 25 µg (n=15), 5 µg (n=15) and 1 µg (n=15) of mRNA-1273. A group of hamsters (n=15) received a prime dose only of 25 µg mRNA-1273 and a mock group received PBS (n=15) at week 0. At week 6, animals were IN challenged with 10^5^ PFU of SARS- CoV-2. On days 2, 4 and 14 post infection, hamsters (n=5 per group) were euthanized for tissue collection. Total serum SARS-CoV-2 S- (B) and RBD-specific (C) IgG titers in serum and (D) neutralizing titers in hamster groups prior to SARS-CoV-2 infection measured by ELISA and plaque reduction assays, respectively. Neutralizing titers were compared to a panel of human convalescent serum samples (Conv.). Bars denote group means ± SE. Significance measured by ANOVA with (B-D) Tukey’s or Sidak’s correction for multiple comparisons between vaccine groups or between time points, respectively (*P ≤ 0.05, **P ≤ 0.01, ***P ≤ 0.001, ****P ≤ 0.0001).

### Prime-boost vaccination effectively protects against SARS-CoV-2 replication in the lungs

Six weeks after prime vaccination (3 weeks after the boost vaccination), all treatment group hamsters were challenged intranasally with 10^5^ plaque forming units (PFU) of SARS-CoV-2. Hamsters were monitored daily for changes in body weight. At 2 and 4 days post infection (dpi), 5 animals from each group were serially euthanized, and the viral load in the right lung and nasal turbinates was determined. The remaining hamsters were observed until the study’s endpoint at day 14. Mock vaccinated hamsters lost an average maximum body weight of 12% by day 6 (Figure 2A and Supplemental Figure 2, A and B). The prime-boost regimen prevented significant weight loss for all but one hamster in the 5 µg prime-boost group; excluding animals necropsied at 2 and 4 dpi, the average maximum weight loss for the combined prime-boost dose vaccine groups was 2.25% over the 14 day infection period (Supplemental Figure 2B). The prime-only vaccinated group lost a maximal mean weight of 6.2%. Moderate inverse correlations were observed between maximum percent weight loss and S-binding IgG and neutralizing antibody titers at week 6 (Supplemental Figure 3A).

**Figure 2.**
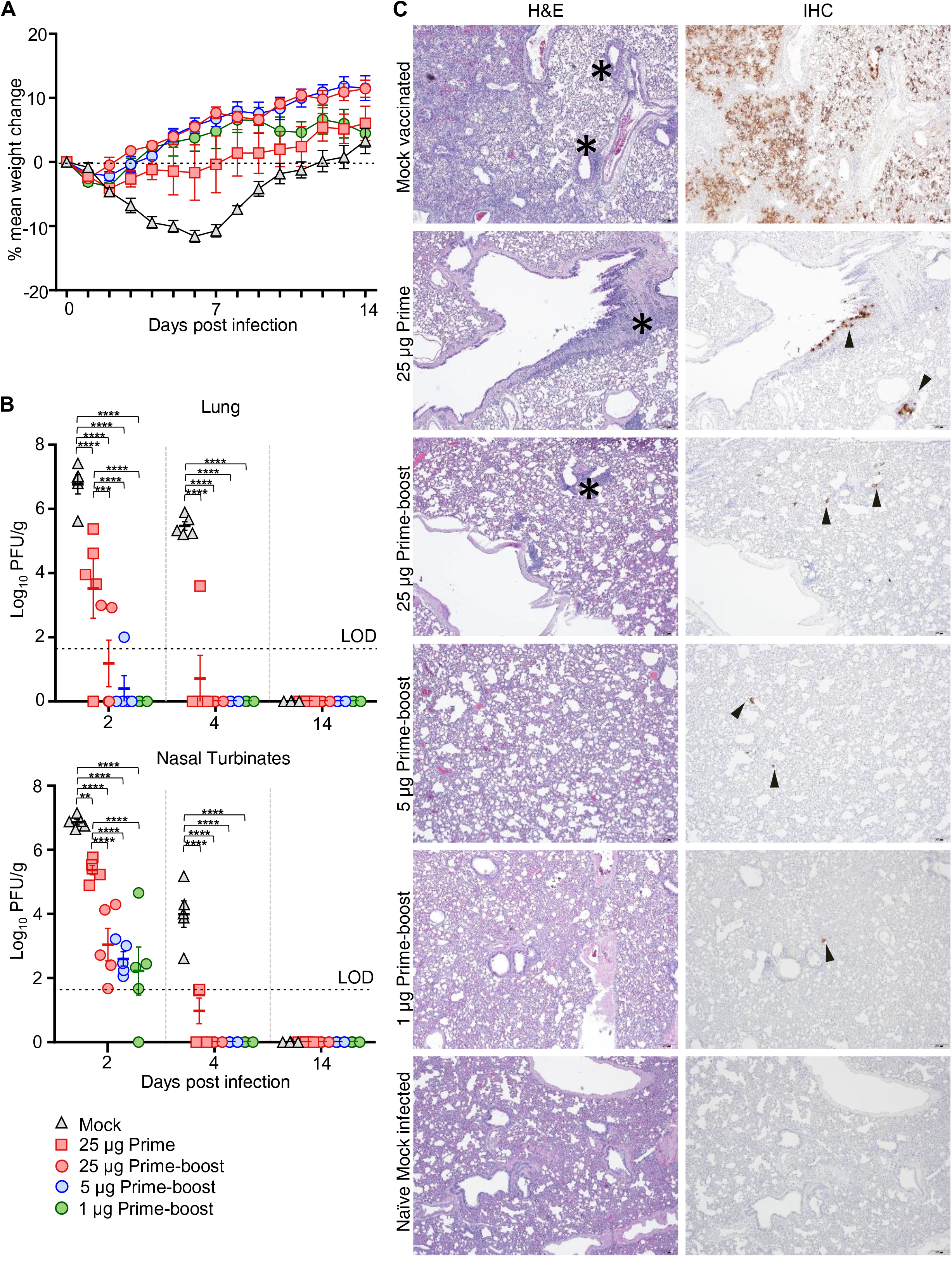
Characteristics of clinical disease following SARS-CoV-2 challenge of vaccinated hamsters. A. Following infection with SARS-CoV-2 six weeks after prime vaccination, hamsters were monitored for mean percent weight change over 14 days after challenge. (B) Viral load expressed in PFU per gram of tissue in the lungs and nasal turbinates at serial endpoint days (2, 4 and 14 dpi). Dotted line indicates limit of detection (LOD). Significance measured by ANOVA with Tukey’s correction for multiple comparisons (*P ≤ 0.05, **P ≤ 0.01, ***P ≤ 0.001, ****P ≤ 0.0001). Error bars represent ± SE. (C) Lung sections from mock vaccinated, prime-only and prime-boost vaccinated groups infected with SARS-CoV-2 and naive mock infected groups at 4 dpi were stained with H&E, and representative photomicrographs (original magnification ×4 (scale bars, 200 µm) as indicated) from each group with virus antigen (arrowhead) in lungs, stained by IHC. * = areas of pv/pbr inflammation (mostly mononuclear).

At 2 dpi, high viral loads were detected by plaque assay in the lungs and nasal turbinates of the mock vaccinated group with a peak mean of 6.8 log_10_ PFU/g and 6.9 log_10_ PFU/g, respectively (Figure 2B). Markedly lower levels of virus were detected in the lungs of all vaccinated groups, while the 1 µg prime-boost dose group had no detectable virus in the lungs. The mean peak virus load in the nasal turbinates were similar among the prime-boost groups and 4.3 log_10_ lower compared to the mock (*P*<0.0001). At 4 dpi, there was no detectable virus in the lungs and nasal turbinates of all prime-boost recipients. Hamsters that received the prime-only vaccine dose had 4.8 log_10_ and 3.0 log_10_ reductions of peak virus load means, respectively, in the lungs and nasal turbinates compared to the control group.

We measured viral subgenomic RNA (sgRNA) in these tissues by qRT-PCR as a potential gauge for replicating virus. Peak sgRNA were detected in the mock group at 2 dpi, with geometric means of 7.4 log_10_ (95% CI range 7.0 log_10_ - 7.9 log_10_) copies/g in the lungs and 7.3 log_10_ (95% CI range 7.1 log_10_ – 7.5 log_10_) copies/g in nasal turbinates (Supplemental Figure 2C). All prime-boost groups showed similar lower sgRNA levels compared to the mock vaccine group with peak geometric means of 2.1 log_10_ (95% CI range 0.86 log_10_ – 3.3 log_10_) sgRNA copies/g in lungs and 6.3 log_10_ (95% CI range 6.4 log_10_ – 6.1 log_10_) sgRNA copies/g in nasal turbinates. In the prime-only vaccine group, sgRNA levels were not significantly reduced in either of these tissues. In close agreement with viral load, no sgRNA was detected in the lungs of prime-boost groups by 4 dpi, except for one hamster in the 1 µg dose group. While viral load was not detected in the nasal turbinates of prime-boost recipients, sgRNA was detected, albeit at markedly reduced levels compared to the mock-vaccinated group. SARS-CoV-2 transmission has been shown to correlate with levels of infectious virus and not sgRNA (22). Therefore, the likelihood of onward transmission was highly reduced by mRNA-1273. The prime-only dose afforded a less robust protection according to sgRNA levels, particularly in the nasal turbinates. Neutralizing antibody titers at week 6 inversely correlated with viral load and sgRNA in both the lungs and nasal turbinates at 2 and 4 dpi (Supplemental Figure 3B and C).

Fourteen days post challenge, the mock vaccinated hamsters had measurable serum S- (Supplemental Figure 4A) and nucleocapsid (NP)-specific (Supplemental Figure 4B) IgG titers and neutralizing titers (Supplemental Figure 4C). An anamnestic response was observed as soon as 4 dpi with the S-specific IgG increasing in recipients of the prime only vaccine regimen. At 14 dpi, all vaccinated hamsters displayed an anamnestic S-specific IgG antibody response. This result contrasts with the unwavering virus-specific IgG levels observed post challenge in vaccinated NHPs (8). Furthermore, NP-specific IgG titers were detected in all groups at 14 dpi (Supplemental Figure 4B) confirming, together with the detection of sgRNA in the upper and lower respiratory tract (Supplemental Figure 2C), replication of the challenge virus before clearance.

### High dose prime-boost vaccination protects against severe pathological changes in the lungs

The lungs of hamsters were evaluated histologically following challenge with SARS-CoV-2 at 2-, 4- and 14-day timepoints (Figure 2C, Supplemental Figure 5 and Supplemental Figure 6). A naïve group of hamsters (*n* = 4), intranasally administered media to mimic virus inoculum and euthanized 4 days later, was included as a control and presented with a moderately prominent alveolar interstitial hypercellularity. While these hypercellular areas may represent regions of atelectasis, the presence of *Pasteurella multocida,* a common respiratory commensal, was also detected by metagenomics analysis of all lung samples used in scRNA-seq (Supplemental Table 1). At 2 dpi, SARS-CoV-2 infection in mock vaccinated animals caused mild interstitial inflammation in the lungs with some animals exhibiting a largely polymorphonuclear cellular infiltrate, comprised predominantly of neutrophils/heterophils, in and around the lung airways (Supplemental Figure 6 and Supplemental Table 2). By 4 dpi, inflammation was largely associated with perivascular and peribronchiolar regions in both a focally diffuse or multifocal distribution affecting, on average, 30-50% of the lung that persisted until the study’s endpoint (Figure 2C, Supplemental Figure 5, Supplemental Table 2). Over the 14-day infection course, prime-boost vaccinated animals displayed mild to moderate inflammation mostly involving the pulmonary interstitium, characterized by expanded alveolar capillary profiles, diminished alveolar spaces and influx of predominantly mononuclear cells (Figure 2C, Supplemental Figure 5, Supplemental Figure 6 and Supplemental Table 2). In the 1 µg prime-boost group, the predominant inflammatory phenotype was mild to moderate interstitial and rarely perivascular- peribronchiolar (4 dpi) inflammation. However, one animal exhibited a more severe pulmonary inflammatory response at 2 dpi, characterized by mild to moderate edema and rare foci of hemorrhage and vascular congestion that was previously described in SARS-CoV-2 infected hamsters (19). Prime-only vaccinated animals generally exhibited reduced inflammation compared to mock-vaccinated animals at 4 dpi; one outlier hamster presented mild pulmonary edema and scant fibrin deposition which was not observed in the rest of the group. Irrespective of vaccination schedule and dose, the airway lumen remained clear of inflammatory cells and cellular debris.

Lung sections were also examined for SARS-CoV-2 antigen at 4 dpi by immunohistochemistry (Figure 2C, Supplemental Figure 5 and Supplemental Table 2). Large amounts of viral antigen were found throughout the lungs of mock-vaccinated hamsters. Prime-boost vaccination limited the amount of detectable total viral antigen more so than prime-only vaccination; SARS-CoV-2 antigen was minimal to absent in the 25 µg and 5 µg prime-boost groups.

Notably, the three outlier hamsters with more severe histopathological phenotypes identified in the 25 µg prime-only group at 4 dpi and in the 1 µg prime-boost group at 2 and 4 dpi, did not exhibit drastic weight loss. Their levels of infectious virus or sgRNA in the lungs or nasal turbinates were also comparable with other group members, with the exception of the single 1 µg recipient at 4 dpi that had detectable sgRNA in the lungs that were not detected in other prime-boost recipients at the same time point. Both 4 dpi outliers had the highest level of viral antigen among all vaccinated hamster samples. Interestingly, all three outliers did not produce neutralizing titers after prime vaccination, a finding that suggests the importance of vaccine dose level and regimen, and the evolving immune response following administration of a second dose.

### Integrated single cell analysis and cell type identification

We performed scRNA-seq on the cranial lung lobe tissue samples from three of the study’s hamster groups that were euthanized at 4 dpi: the naive hamsters mock-infected with media (Naive, N, *n* = 4) and the 5 µg prime-boost vaccinated (Vaccinated + Infected, VI, *n* = 5) and mock-vaccinated (Mock-Vaccinated + Infected, MI, *n* = 5) groups infected with SARS-CoV-2. Lung samples from the N group served as a baseline control for analysis. The scRNA-seq data were aligned against the hamster and SARS-CoV-2 genomes and integrated using the Seurat scRNA-seq analysis pipeline (24). The expression profile of one VI animal that showed excessive weight loss at 4 dpi (Supplemental Figure 2A and Supplemental Figure 7A) was very distinct, so it was analyzed separately and not included in the integrated analysis. Integration of the data from the remaining 13 samples identified 15 cell types, including multiple T cell, dendritic cell (DC) and macrophage subtypes (Figure 3A). The main markers that distinguish the cell types are shown in Supplemental Figure 8.

**Figure 3.**
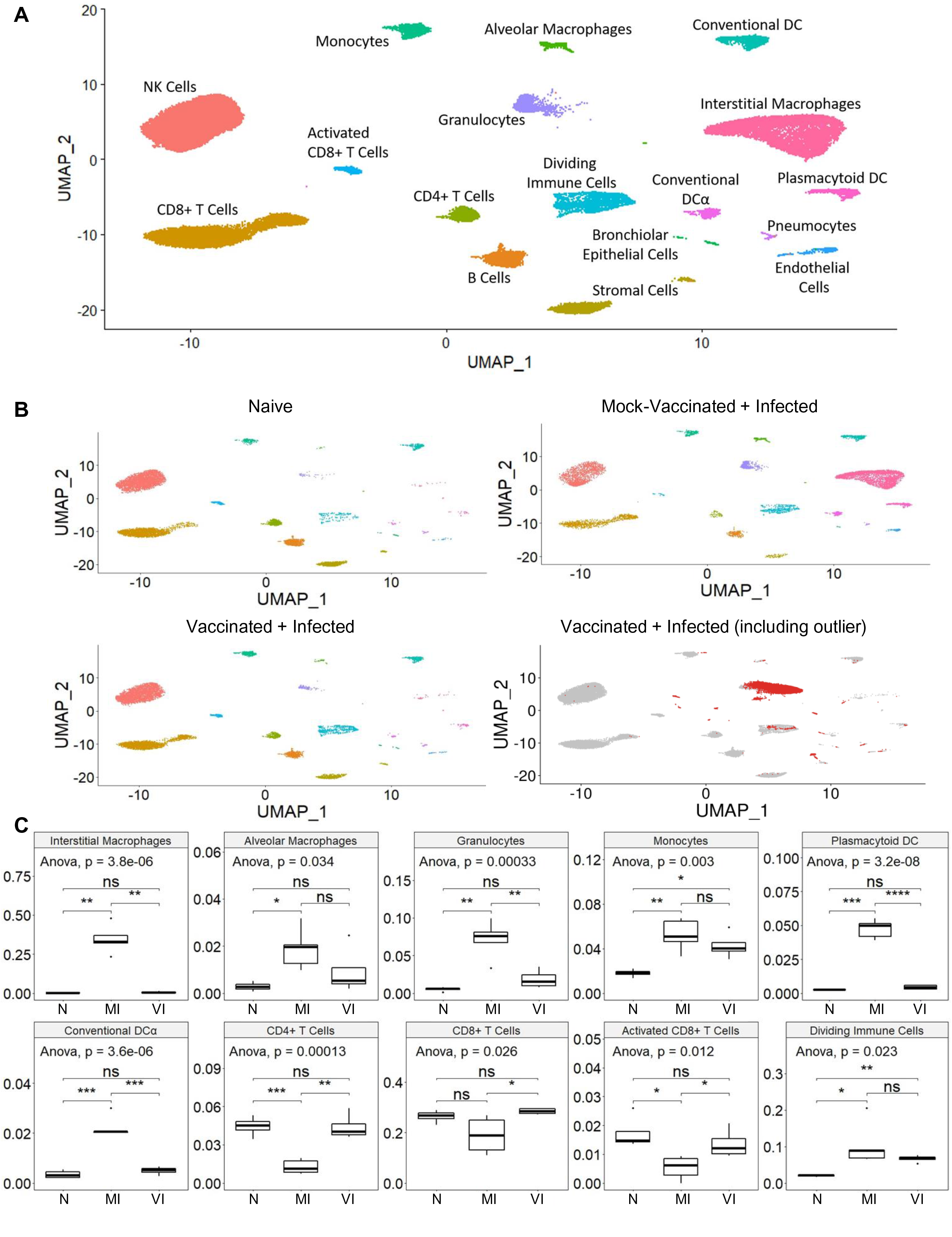
Single-cell RNA-seq analysis identifies cell population differences among Naive (N), Mock-vaccinated Infected (MI) and Vaccinated Infected (VI) pulmonary tissue. (A) Integrated UMAP of all N, MI and VI samples excluding one VI outlier sample. (B) UMAP of the N, MI and VI groups and VI outlier. N, MI and VI plots were each randomly downsampled to 8,000 cells to facilitate comparison. Cell identities correspond to panel A. The VI outlier sample is red and the other VI samples are shown in grey for reference. (C) Box plots comparing cell-type proportions observed in each group. ANOVA and post-hoc Tukey test pairwise comparisons are shown. * p<.05, ** p<.01, ***p<.001, **** p<.0001.

Visualizing these 13 datasets by group on the integrated UMAP coordinates showed an overall change in cell type proportion across MI and VI groups when compared to the N group. Most noticeably, there was a more than 30% increase in the presence of interstitial macrophages in the MI hamsters compared to a negligible fraction in N hamsters (Figure 3, B and C). Importantly, this increase in macrophages was not seen in VI hamsters. Significant increases in proportion of alveolar macrophages, DC subtypes (plasmacytoid DC (pDC), conventional DC (cDC) and activated DC (DC α)), and granulocytes were also found in MI hamsters and not VI hamsters. We also detected decreases in proportion of CD4^+^, CD8^+^ and CD8^+^ activated T cells in MI hamsters and not VI hamsters. The MI and VI hamsters both showed increases in the proportion of the dividing immune cells and monocytes when compared to N hamsters. The moderate changes in the lung cell composition of VI hamsters and infiltration of inflammatory cells in lungs of MI hamsters are consistent with histology results described in the previous section.

Overlaying the data from the outlying VI hamster onto the UMAP coordinates, together with the other 4 VI samples, showed different cell type composition in this sample (Figure 3B lower right). Granulocytes represented over 75% of the cells in this outlier sample but were less than 5% of cells in the other VI samples. The gene expression pattern in the granulocytes from the outlier VI sample also differed from the granulocytes in the other VI samples (Supplemental Figure 7, B and C). Pathway analysis of the differentially expressed genes (DEGs) in granulocytes from the outlier showed upregulation of classical neutrophil processes associated with degranulation, activation and proliferation, and downregulation of immune suppressive interleukins (Supplemental Figure 7D). The abnormal cellular composition in the VI outlier was likely due to sampling of heterophilic cellular lesions constrained to the cranial lobe given histopathological analysis on the remaining left lung showed drastic reduction of inflammation. The activity of heterophils concentrated at a lesion is expected to be different to those patrolling the lung parenchyma, adopting a functional state at the site of a foreign antigen.

### Vaccination attenuates infection of immune cells and their activation

The scRNA-seq reads were mapped to the SARS-CoV-2 genome to identify which cell types contained viral RNA. Viral gene reads were detected only in MI hamsters (Figure 4, A-C), consistent with the clearance of infection in the lungs of the VI hamsters that was observed by qRT-PCR (Supplemental Figure 2C). Several MI samples with higher overall viral reads had virus gene expression detected in a broad selection of cell types in the lungs, including lymphoid and myeloid cells (Figure 4A). Macrophages and granulocytes had the highest average viral read counts. Viral ORF1ab was the highest expressed viral gene (Supplemental Figure 9).

**Figure 4.**
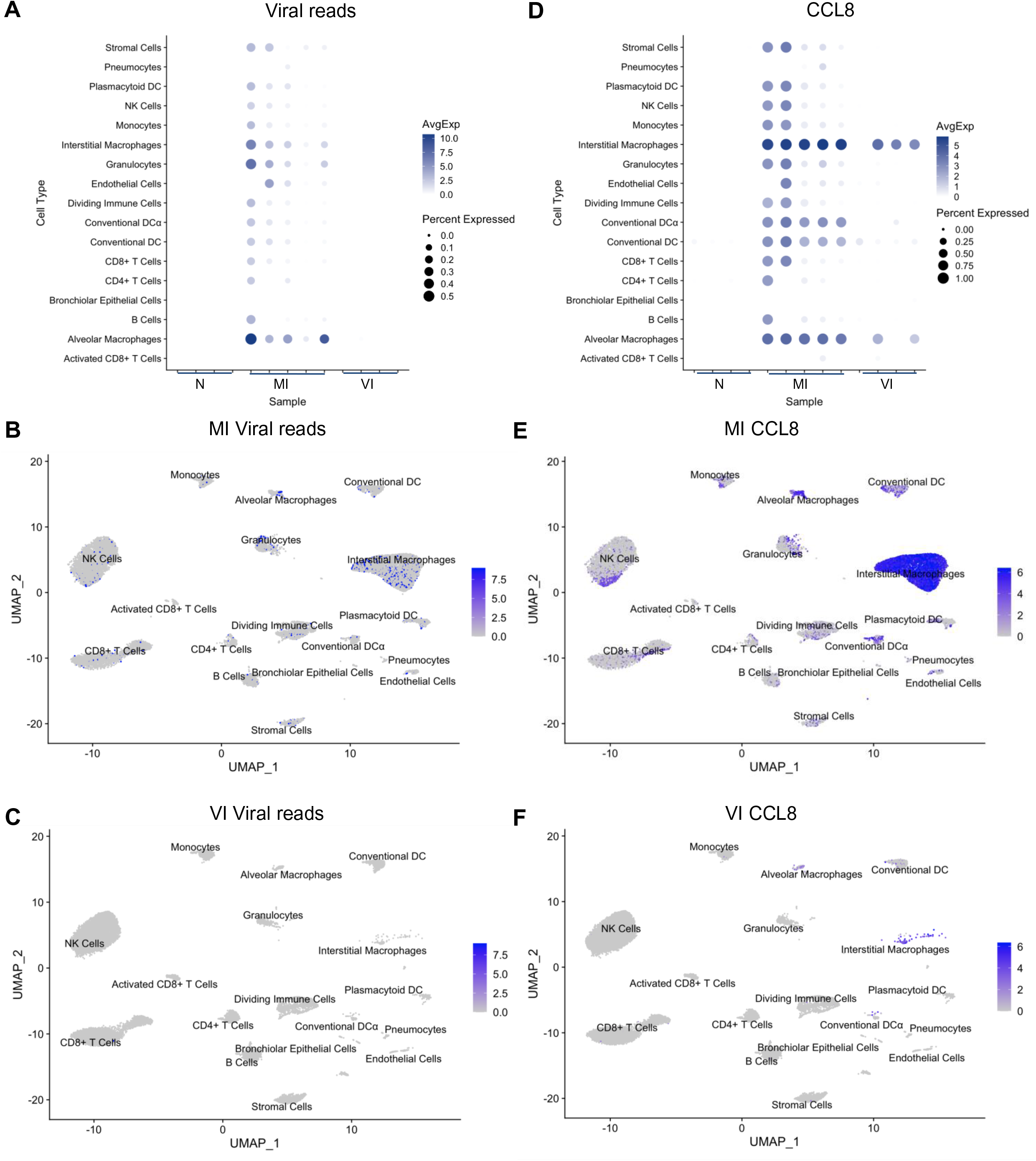
Viral reads and CCL8 expression from lung tissue single cell RNA-seq. (A) Dot plot showing expression of all viral reads in each cell-type in each sample. (B-C) Per-cell expression levels of all viral reads in (B) Mock-Vaccinated + Infected and (C) Vaccinated + Infected visualized in UMAP plot. (D) Dot plot showing expression of CCL8 in each cell-type in each sample. (E-F) Per-cell expression levels of CCL8 in (E) Mock-Vaccinated + Infected and (F) Vaccinated + Infected visualized in UMAP plot.

Expression of CCL8, the chemoattractant for monocytes and macrophages reported to be significantly induced in response to SARS-CoV-2 infection in ferret upper respiratory tract (17), was minimal in the lungs of N animals but most up-regulated in the MI hamsters (Figure 4, D and E). The upregulation of CCL8 is consistent with macrophage infiltration of the lung observed in the MI group and in COVID-19 patients (25). The cellular distribution of CCL8 expression in MI lung mirrored that of the virus RNA, being highest in the interstitial and alveolar macrophage populations. In the VI group, increases in CCL8 expression was only observed in the macrophage populations in some of the samples studied, despite the absence of SARS-CoV-2 mRNA (Figure 4, D and F). Overall, these data suggest that vaccination impedes SARS-CoV-2 infection of lymphoid and myeloid cells and reduces their activation.

### Following SARS-CoV-2 infection, similar transcriptional programs are regulated in lung immune cells from vaccinated and mock-vaccinated hamsters

To determine whether immune cells in the lungs showed differences in their transcriptional states in the three groups studied, we focused on two lymphoid cell types critical for viral clearance during respiratory infections, CD8^+^ T cells and NK cells, which were relatively abundant across all samples. We also examined myeloid cells, including pDCs, cDCs, and monocytes, which bridge innate sensing with adaptive responses. We performed differential expression analysis for each cell type (see Methods), comparing the MI to N samples and the VI to N samples. This analysis in CD8^+^ T cells showed that less transcripts were differentially expressed in the CD8^+^ T cells from VI lungs compared to N than CD8^+^ T cells from MI lungs compared to N, with the DEGs in VI lungs being predominantly a subset of those regulated in the cells from MI samples (Figure 5A). A scatter plot comparing the log fold-changes of the set of transcripts significantly regulated (FDR <0.05) in either MI or VI comparisons to N showed that changes of these transcripts were highly correlated (p<2.2e^∧^-16). The slope of the linear regression fit was less than 1, indicating that the gene expression changes were relatively lower in the VI samples. These results suggested that similar transcriptional programs were modulated in these immune cells in the MI and VI hamsters, although the regulation was to a lesser extent in the VI hamsters. Similar analyses of the NK cells, pDC, cDC, and monocytes revealed that the expression of DEGs for both MI and VI conditions compared to N was also highly significantly correlated (Figure 5B, Figure 6, A and B and Supplemental Figure 12A). The slopes of the regression lines showed that the changes were also of lower magnitude in the VI-derived cells. Overall, these results indicated that across all cell types analyzed, the transcriptome regulation in the VI group was similar to that of MI group, but the changes were of a lower magnitude in the VI group.

**Figure 5.**
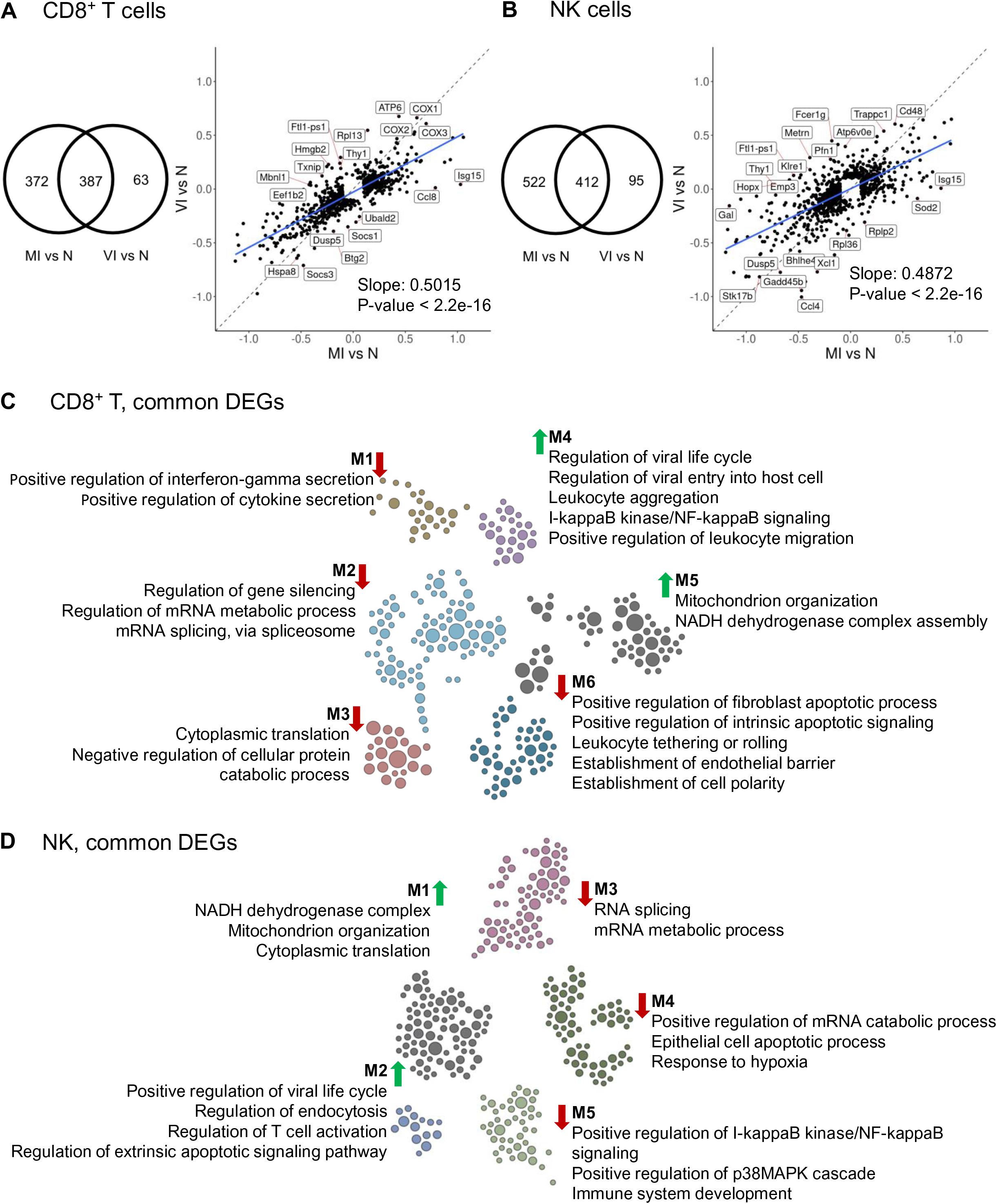
Functional pathway analysis of MI vs. N and VI vs. N comparisons in CD8^+^ T cells and NK cells. (A) Venn diagram of the union of DEGs of the two comparisons in CD8^+^ T cells. Scatter plot of the logFC of the union of DEGs of the two comparisons in CD8^+^ T cells, i.e. the same group of genes as in the Venn diagram. For each comparison, DEGs were selected with thresholds FDR < 0.05 and absolute logFC > 0.1. Linear regression model was fitted to the scatter plot (adjusted R^∧^2 = 0.6687). (B) Venn diagram of the union of DEGs of the two comparisons in NK cells. Scatter plot of the logFC of the union of DEGs of the two comparisons in NK cells, i.e. the same group of genes as in the Venn diagram. For each comparison, DEGs were selected with thresholds FDR < 0.05 and absolute logFC > 0.1. Linear regression model was fitted to the scatter plot (adjusted R^∧^2 = 0.5147). (C,D) Functional network showing modules of enriched pathways using DEGs shared between two comparisons in (C) CD8^+^ T cells and (D) NK cells. Each gene is represented by a circle and color coded according to module. The size of each circle reflects its connectivity in the network. Edges are not shown, allowing for easy viewing. The module label is shown with the functional processes and pathways identified in each module. Up or down regulation of pathway is indicated by arrow direction.

**Figure 6.**
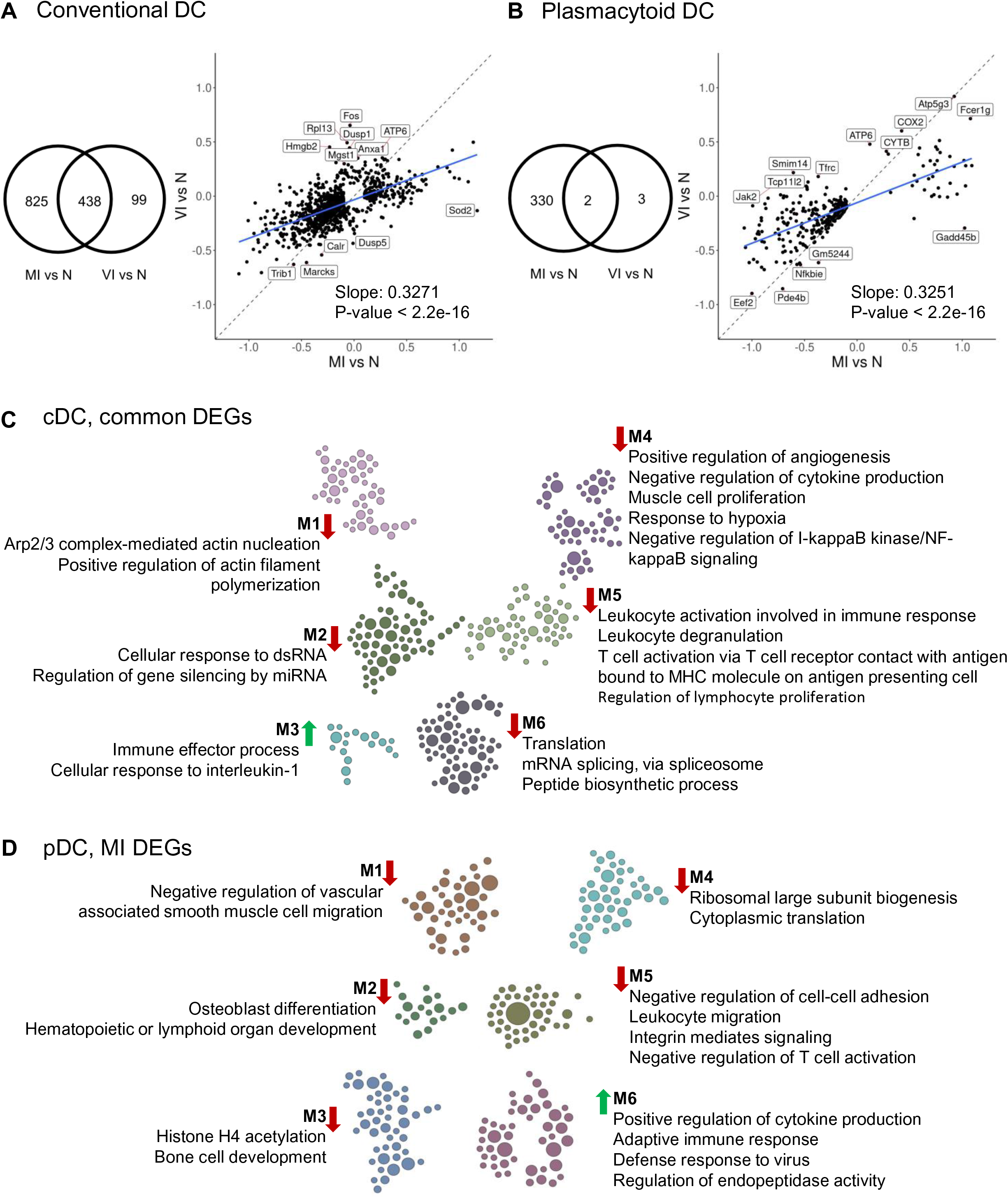
Functional pathway analysis of MI vs. N and VI vs. N comparisons in conventional DC (cDC) and plasmacytoid DC (pDC). (A) Venn diagram of the union of DEGs of the two comparisons in cDC. Scatter plot of the logFC of the union of DEGs of the two comparisons in cDC, i.e. the same group of genes as in the Venn diagram. For each comparison, DEGs were selected with thresholds FDR < 0.05 and absolute logFC > 0.1. Linear regression model was fitted to the scatter plot (adjusted R^∧^2 = 0.4042). (B) Venn diagram of the union of DEGs of the two comparisons in pDC. Scatter plot of the logFC of the union of DEGs of the two comparisons in pDC cells, i.e. the same group of genes as in the Venn diagram. For each comparison, DEGs were selected with thresholds FDR < 0.05 and absolute logFC > 0.1. Linear regression model was fitted to the scatter plot (adjusted R^∧^2 = 0.5098). (C,D) Functional network showing modules of enriched pathways using (C) DEGs shared between two comparisons in cDC or (D) DEGs unique to MI pDC. Each gene is represented by a circle and color coded according to module. The size of each circle reflects its connectivity in the network . Edges are not shown, allowing for easy viewing. The module label is shown with the functional processes and pathways identified in each module. Up or down regulation of pathway is indicated by arrow direction.

To elucidate the common pathways regulated in the MI and VI groups, we performed a module-based functional enrichment analysis. Among the modules within the set of upregulated DEGs, immune activation and viral response were noted across CD8^+^ T cells, NK cells, cDC, and monocytes (Figure 5, C and D, Figure 6C and Supplemental Figure 12B). Common pathway enrichment analysis could not be performed for the pDCs due to limited DEGs identified in the VI group.

Since we generally saw a lower degree of gene activation in the VI group, less genes passed the statistical threshold for differential expression in that group as compared to the MI group. Given the general conservation of the transcriptional programs between the MI and VI groups, a substantial number of DEGs were common to the two comparisons while another large number of DEGs were specific to the MI group, and relatively few DEGs were specific to the VI group (see Venn diagrams in Figure 5, A and B, Figure 6A and Supplemental Figure 12A). We performed module-based functional enrichment analysis on the DEGs that were specific to either MI or VI groups for each cell type analyzed.

Similar pathways were modulated in MI-specific NK DEGs and CD8^+^ T cell DEGs; viral response, migration, regulation of apoptotic signaling, and cellular responses to interferon-γ and oxidative stress were upregulated, while homeostasis and cellular maintenance were downregulated (Supplemental Figure 10A and Supplemental Figure 11A). However, the proliferation process was upregulated in CD8^+^ T cells and negatively regulated in NK cells. Type I interferon production and viral response pathways were upregulated, and cellular maintenance pathways were downregulated in MI-specific cDC DEGs and monocyte DEGs (Supplemental Figure 12C and Supplemental Figure 13A). Adaptive immune response, including T-cell activation, was modulated in both MI-specific pDC DEGs and MI-specific cDC DEGs (Figure 6D and Supplemental Figure 13A). Finally, additional viral response pathways were upregulated in MI-specific pDC DEGs (Figure 6D). Since fewer DEGs were identified as VI-specific, there were few regulated pathway modules detected (Supplemental Figure 10B, Supplemental Figure 11B and Supplemental Figure 13B).

Overall, these data show that SARS-CoV-2 infection induced largely similar transcriptional programs in immune cells for both VI and MI animals, including many viral response and immune activation pathways. In MI-specific transcriptional programs, positive regulation of cytokine signaling stood out as enriched across all cell types studied, while cellular maintenance functions were typically downregulated. On the other hand, in the VI animals, the magnitude of gene expression was lower and fewer differential genes were detected, resulting in few regulated pathway modules.

## DISCUSSION

In humans, COVID-19 can progress to severe clinical disease which manifests as pneumonia. mRNA-1273 was previously shown to be efficacious in NHPs (8), mice (1) and recently, in a large Phase 3 trial (11). Hamsters consistently exhibit the hallmarks of severe COVID-19 disease and are therefore an important model for preclinical vaccine efficacy studies. We show that two doses of mRNA-1273 reduced viral load in the upper and lower airways of hamsters and protected against weight loss while a prime-only vaccination provided partial protection. Two doses of mRNA-1273 were required to induce neutralizing titers comparable with the higher titers seen in convalescent COVID-19 patients. Although neutralizing titers were not dependent on dosage in a prime-boost schedule, the highest prime-boost dose of 25 µg provided better protection against lung injury and weight loss. Two hamsters which experienced more severe weight loss were identified in the low prime-boost dose or prime-only vaccine groups despite having a high binding and neutralizing antibody titers. Moreover, despite a strong inverse correlation between neutralizing antibody titers and virus load in the respiratory tract, the inverse correlation between neutralizing or binding antibody titers and weight loss was only moderate. Other antibody- and cell-mediated mechanisms may therefore be required for complete protection against SARS-CoV-2 disease in this model. This emphasizes the importance of a qualitative, not just quantitative, immune response, which may depend on the vaccine dose and regimen. The efficacy of mRNA-1273 in hamsters is distinguished from efficacy studies with NHP and mouse models, as protection was afforded against more severe pathological phenotypes and clinical disease, and the immunity was non-sterilizing.

Cellular responses in vaccinated hamsters appeared to promote pulmonary homeostasis and a less proliferative and migratory response following infection while supporting virus clearance. While enriched gene expression pathways associated with immune effector functions were shared by both VI and MI hamsters, pathways regulating cell migration were generally unique to MI hamsters. Granulocytes and inflammatory macrophages were abundant in the lungs of MI hamsters, features that are also observed in patients with severe COVID-19 (25, 26). High viral RNA reads within these cells suggests acquisition of infected cells by phagocytosis or a susceptibility to infection. Vaccination averted pro-inflammatory responses and the influx of these innate immune cells. The dominant CCL8 response across all immune cells in MI hamsters was restricted to macrophages in VI hamsters and may represent a signal for regulated cellular trafficking as opposed to inflammation. Similar levels of monocytes infiltrated the lungs of both MI and VI hamsters. While monocytes from both MI and VI hamsters possessed DEGs compared to N samples that modulated their responses to external stimulus, the monocytes from MI, but not VI, hamsters exhibited upregulated inflammatory characteristics that are observed in severe COVID-19 cases (27). The controlled responses in VI monocytes suggest a supportive role in immune regulation. Upregulated adaptation to oxidative stress in monocytes, NK cells and CD8^+^ T cells was unique to MI hamsters, indicating that vaccination controlled exacerbated responses to infection which promotes tissue damage and hypoxic respiratory failure in severe patients (28). Certain NK cell phenotypes were associated with COVID-19 related hyperinflammation and similar to the populations in the bronchoalveolar lavage fluid (BALF) from patients with severe disease (29); the cytotoxic functions of NK cells in MI hamsters were enriched while their proliferation processes were negatively regulated. NK cells in VI hamsters were functioning in a similar albeit attenuated capacity, to those in MI hamsters, which suggests a reduced ability to clear virus-infected cells. NK cells maintained immune homeostasis in the lungs upon infection of mice vaccinated against influenza, rather than realizing their killing capacity, to enable virus clearance by the adaptive immune system with minimal pathology (30). Thus mRNA-1273 vaccination prevented the hyperactivation of the innate immune system following infection but permitted a controlled cellular antiviral response conducive to preserving the lung milieu.

Patients with severe COVID-19 display aberrant T cell activation and differentiation, lymphopenia (31, 32), and generally have more proliferative T cells but less CD8^+^ T cell proportions with limited clonal diversity in BALF (25). Conversely, non-hospitalized, recovered individuals have virus-specific T cell memory (33, 34). In the lungs of MI hamsters, T cell proportions declined indicating that extravasation to affected tissues, one of the suggested causes of lymphopenia in severe patients, was not occurring. Instead, macroautophagy, a process which supports cell survival against environmental stresses or upon the activation of naïve T cells (35, 36), was downregulated in CD8^+^ T cells of MI hamsters despite their responsiveness to oxidative stress and cytokine stimulus. Together with an extended lifespan, reduced telomerase maintenance and upregulated proapoptotic pathways, activated CD8^+^ T cells in MI hamsters may develop an effector phenotype but are replicative senescent, susceptible to depletion or unable to form functional memory T cells (37, 38). CD8^+^ T cell exhaustion from hyperactivation or augmented expression of pro-apoptotic molecules has been linked to their depletion in severe COVID-19 cases (39, 40). Vaccination prevented a reduction in T-cell frequencies in the lungs. Moreover, CD8^+^ T cells in VI hamsters were less enriched for activation and proliferative pathways compared to MI hamsters, but sustained effector functions. This suggests the CD8^+^ T cell response in VI hamsters was either from memory effectors established after vaccination, to facilitate viral clearance, or from clonally diverse, tissue-resident cells with an effector phenotype after SARS-CoV-2 challenge, as seen in the BALF from moderate but not severe COVID-19 patients (25).

Aberrant T cell responses or their depletion in severe COVID-19 may be indirectly attributed to DCs that are impaired in maturation and T cell activation (41). DC subsets infiltrated the lungs of MI hamsters but remained unaltered in VI hamsters. Similar to acute phase coronavirus infections (42), infiltrating pDCs may provide rapid antiviral responses through the upregulation of cytokine production pathways. However, pDCs were not necessary for influenza A virus clearance and were shown to have a deleterious role on CD8^+^ T cells during lethal infection of mice (43, 44). In MI hamsters, the lymphoid organ development pathway of pDCs was downregulated suggesting their role in T cell differentiation was compromised. The low numbers of pDCs in VI hamsters and their limited DEGs suggests they were not involved in vaccine-mediated responses to infection. Dysfunctional T cell activation in COVID-19 patients has been associated with the downregulation of MHC and costimulatory molecules on antigen presenting cells (41, 45, 46). While immune effector pathways and chemotactic cues from IL-1 to migrate to the lung-draining lymph nodes were upregulated in cDCs from both MI and VI hamsters, DEGs involved in the activation of T cells via MHC molecules were paradoxically downregulated. Despite this remarkable similarity in MI and VI-induced cDC pathways, disparities in DEG expression magnitude, cDC numbers and external stimulus from the lung milieu may account for different cDC-mediated outcomes. Diminished MHC-dependent T cell activation by cDCs from MI hamsters likely impaired the adaptive immune response, while in VI hamsters, the controlled cell contact dependent activation by cDCs may be sufficient for recall responses or sustaining the effector functions of CD8^+^ T cells (47). mRNA-1273 promoted a controlled response to SARS-CoV-2 infection that prevented desynchrony between the innate and adaptive immune arms which can exacerbate inflammation and disease severity.

The incomplete annotation of the hamster genome prevented identification of highly specific subtypes within immune cell populations. Cell type identification partially based on mouse data and the pathway analysis based on human gene ontology terms did permit detailed profiling of cellular responses. While this study examined the response to infection directly in the lungs of VI and MI hamsters rather than BALF, scRNA-seq analysis was limited by the fact that one region of lung was studied whereas responses to an infection within the lungs may be spatially distinct. As a result, we captured a unique perspective on a potentially focal inflammatory lesion comprised of activated neutrophils/heterophils formed after infection of a low vaccine-dose VI hamster. Nevertheless, we excluded this scRNA-seq outlier to effectively illustrate the cellular response dynamics at the site of infection in vaccinated and mock-vaccinated hamsters exposed to SARS-CoV-2.

mRNA-1273 provided hamsters with infection-permissive immunity that following SARS-CoV-2 infection reduced virus load and severe disease by promoting humoral responses in the periphery and antiviral cellular responses in the lung. mRNA-1273 demonstrated protection is predominately achieved through the induction of antibodies, and cellular responses may act to support complete clearance of the virus. Importantly, the safety of mRNA-1273 was indicated by the absence of aberrant cellular pathways in VI hamsters after challenge. Transient, low level virus replication in VI hamsters did trigger the regulation of cellular programs in some immune cells that were strikingly similar to MI hamsters. However, the responses were attenuated and lacked the atypical manifestations which contribute to inflammation and lung injury in critical COVID-19 cases. The surprising commonalities in transcriptional pathway regulation between vaccinated and mock vaccinated recipients during acute infection warrants investigation into how immunity induced by other vaccines is regulated following infection to provide the foundation for successful vaccination.

## METHODS

### Pre-clinical mRNA-1273 mRNA and LNP production process

Pre-clinical mRNA-1273 is a purified mRNA transcript encoding the prefusion-stabilized SARS-CoV-2 S-2P protein and encapsulated by a lipid nanoparticle. The process for mRNA synthesis, purification, and encapsulation was described previously (1).

### Human convalescent-phase serum

Human convalescent sera (n = 6) were obtained from adults between 18 and 55 years old with mild (*n* = 2), medium (*n* = 2), and severe (n = 2) COVID-19 and a history of laboratory-confirmed SARS-CoV-2 infection 1 to 2 months before providing specimens. In addition, SARS-CoV-2 naïve sera samples (*n* = 3) were also included in analyses. These specimens were obtained from Aalto Bio Reagents Ltd.

### ELISA

S, RBD or nucleocapsid proteins (1µg/mL, Sino Biological) were coated onto 96-well plates for 16 h. Plates were then blocked with SuperBlock (Pierce). Five-fold serial dilutions of hamster serum were then added to the plates (assay diluent – PBS + 0.05% Tween-20 + 5% goat serum) and incubated for 2 hours at 37°C. Bound antibodies were detected with HRP-conjugated goat anti-hamster IgG (1:10,000 Abcam AB7146). Following the addition of TMB substrate (SeraCare #5120-0077) and TMB stop solution (SeraCare #5150-0021), the absorbance was measured at OD 450 nm. Titers were determined using a four-parameter logistic curve fit in GraphPad Prism (GraphPad Software, Inc.) and defined as the reciprocal dilution at approximately OD450nm = 1.5 (normalized to a hamster standard on each plate).

### SARS-CoV-2 neutralization assay

Two-fold serial dilutions of heat-inactivated serum at an initial dilution of 1:10 were prepared in serum-free MEM media and incubated with SARS-CoV-2-mNG for 1 hour at 37°C at a final concentration of 100 PFU. Virus-serum mixtures then were absorbed onto Vero-E6 monolayers in black optical 96 well plates for 1 hour at 37°C and replaced with MEM/Methylcellulose/2% FBS overlay. After 2 days of incubation at 37°C in humidified 5% CO2, neon green plaques were visualized and counted. Neutralization titers at an end point of 60% plaque reduction were determined.

### Hamster study

Three groups of 6-7 week female golden Syrian hamsters (Envigo) (*n* = 15 per group) were vaccinated with mRNA-1273 diluted to 25 µg, 5 µg and 1 µg in PBS for prime-boost vaccine regimens. An additional group (*n* = 15) was prime vaccinated only with 25 µg of mRNA-1273. A mock vaccinated control group (*n* = 15) received PBS only. Formulations were administered by IM injection to each hind leg (50 µL per dose site). At week 3, prime-boost groups received their second vaccine dose. At week 6 (day 42), all animals were infected via the intranasal route with 100 µL of 2019-nCoV/USA-WA01/2020 (GenBank: MN985325.1, courtesy World Reference Center for Emerging Viruses and Arboviruses, the University of Texas Medical Branch (UTMB)) at 10^5^ PFU (approximately 50 µL per nostril). Over the course of the study, hamsters were monitored daily for clinically for weight changes. On each serial endpoint day (days 2, 4 and 14 post-challenge), lungs, nasal turbinates and serum were collected from 5 hamsters per group. All animal protocols were approved by the Institutional Animal Care and Use Committee at UTMB.

### Analysis of viral load by plaque assay

The right lung and nasal turbinates were homogenized in Leibovitz L-15 medium (Thermo Fisher Scientific, Cat No. 11415064)/10% FBS/1X Antibiotic-Antimycotic (Thermo Fisher Scientific, Cat No.15240062) using the TissueLyser II bead mill (Qiagen) and 5 mm stainless steel beads (Qiagen, Cat No. 69989) and briefly centrifuged. Ten-fold serial dilutions of homogenates were prepared in serum free MEM media and absorbed on Vero-E6 monolayers in 48 well plate for 1 hour at 37°C. The virus inoculum was removed replaced with an overlay of MEM/methylcellulose/2% FBS. After 3 days, plaques were immunostained with a human monoclonal antibody cocktail specific for the S protein (Distributed Bio) and an anti-human IgG HRP conjugated secondary antibody (Sera Care Cat No.5220-0456). Plaques were counted and virus load per gram tissue was determined.

### Analysis of viral load by qRT-PCR

Replicating viral RNA was determined in the lungs and nasal turbinates by measuring subgenomic SARS-CoV-2 E gene RNA by qRT-PCR using previously described primers, probe and cycle conditions (48). Briefly, RNA was extracted from lung and nasal turbinates homogenates using TRIZOL LS (Thermo Fisher Scientific) and Direct-zol RNA Microprep kit (Zymo Research). RNA (500 ng) was reverse transcribed using Superscript IV (Thermo Fisher Scientific) according to the manufacturer’s recommendations. Quantitative real-time PCR was perform using the TaqMan Fast Advanced Master Mix (Thermo Fisher Scientific), and primers and a FAM-ZEN/Iowa Black FQ labeled probe sequence (IDT) on the QuantStudio 6 system (Applied Biosystems). An Ultramer DNA oligo (IDT) spanning the amplicon was used to generate standard curves to calculate the sgRNA copies per gram of tissue.

### Histopathology

Lung samples were processed per a standard protocol for histological analysis. Briefly, the lower left lung lobe was fixed in 10% neutral buffered formalin, embedded in paraffin, sectioned at 5 μm and stained with Hematoxylin and Eosin (H&E) for routine histopathology. Samples were evaluated by a board-certified veterinary pathologist in a blinded manner. Sections were examined under light microscopy using an Olympus BX51 microscope and photographs were taken using an Olympus DP73 camera. Representative sections were displayed.

### Immunohistochemistry

Staining of hamster lung sections was done using the Bond RX automated system with the Polymer Define Detection System (Leica) as per manufacturer’s protocol. Tissue sections were dewaxed with Bond Dewaxing Solution (Leica) at 72°C for 30 min then subsequently rehydrated with graded alcohol washes and 1x Immuno Wash (StatLab). Heat-induced epitope retrieval (HIER) was performed using Epitope Retrieval Solution 1 (Leica), heated to 100°C for 20 min. A peroxide block (Leica) was applied for 5 min to quench endogenous peroxidase activity prior to applying the SARS-CoV-2 antibody (1:2000, GeneTex, GTX135357). Antibodies were diluted in Background Reducing Antibody Diluent (Agilent). The tissue was subsequently incubated with an anti-rabbit HRP polymer (Leica) and colorized with 3,3’-Diaminobenzidine (DAB) chromogen for 10 min. Slides were counterstained with hematoxylin and representative sections were shown.

### Single cell isolation and RNA sequencing

The cranial left lung sections taken at the time of necropsy at 4 dpi from the 5 µg prime-boost group (VI, *n* = 5), mock vaccinated group (MI, *n* = 5) and a naïve group (N, *n* = 4) of hamsters that were mock infected via the IN route with 100 µL of media inoculum. Lung samples were enzymatically digested and homogenized using the Lung Dissociation kit, mouse (Miltenyi Biotec Cat No. 130-095-927) and GentleMACS Dissociator (Miltenyi Biotec) according to the manufacturer’s protocol. After 30 min of digestion, the samples were filtered through a 70 µM filter, and RBC lysis was performed (ThermoFisher, Cat #00-4333-57). After two washes in PBS containing 0.05 mM EDTA and 0.04% BSA, the cell pellet was re-suspended in 1ml of buffer and filtered through a 40 µM Flowmi Cell strainer (Bel-Art # H13680-0040). The cell viability and concentration were determined on a TC20 cell counter (Bio-Rad, CA). Dead cell removal was performed if the viability was lower than 80% using Dead Cell Removal Kit (Miltenyi Biotec, Cat#130-090-101) as per manufacturer’s recommendations.

Seven thousand cells were targeted for generation of barcoded gel bead emulsions using the Chromium Single Cell 3’ version 3.0 chemistry (10x Genomics, Cat# 1000077). After reverse transcription, the cDNA was amplified and purified using SPRISelect magnetic beads (Beckman Coulter, CA, Cat#B23317). The purified cDNA was precipitated in 80% ethanol, removed from BSL-4 containment, tested for quality on Bioanalyzer, and 3’ gene expression libraries were prepared as per manufacturer’s instructions. Libraries were quantified, and pooled libraries were submitted for sequencing (∼140,000 reads per sample) on Novaseq S1 Flow cell (New York Genome Center).

### Processing of scRNA-seq data

scRNA-seq data were processed using the Cell Ranger pipeline (v4.0.0). Alignment was done against the Mesocricetus auratus Ensembl genome (MesAur1.0). STARsolo was used to quantify reads associated with the viral genome by aligning against the SARS-CoV-2 genome (ASM985889v3). The 5’ leader sequence and 3’ UTR of the SARS-CoV-2 genome were included as reference gtf entries. Resulting count tables were then analyzed using Seurat (V3.1.5) (49). Cells with abnormally high mitochondrial gene content (greater than 20% of UMIs per cell) and/or cells containing less than 300 unique genes were removed. A single sample in the VI group was a clear outlier (see Supplemental Figure 7) and was removed from all further analysis. For each group (MI: *n* = 5, VI: *n* = 4, and N: *n* = 4), all samples were merged in an unbiased manner. The three resulting datasets (one per group) were then integrated using the “IntegrateData()” function of Seurat, to account for potential batch effects. Finally, the PCA and UMAP (50) projections were calculated for the integrated sample set using the top 2000 highly variable genes, as determined by Seurat, and significant cell clusters were identified.

### Cell type identification for scRNA-seq data

Integration of the data from 13 samples comprising 50,694 cells identified 17 cell types. Single-cell Mouse Cell Atlas (scMCA) (51) was used to guide cell type identification. Reference gene markers were initially determined from the MCA adult lung dataset. For each cell-type, marker genes were defined as genes satisfying two conditions. Then, the average logFC of each marker gene was greater than 1.0 as compared to the other cell types. Second, the fraction of cells expressing the marker gene in the cell type was higher than the fraction of cells expressing the marker gene in all other cell types by at least 0.4. Using the defined marker gene lists, the correlation of the average gene expression in each cell type in the MCA adult lung was computed with each cell in our dataset, annotating each cell with the most highly correlated cell type. Additional marker sets and prior knowledge (52) was used for some cell-types that were ambiguous or missing from the reference data (granulocytes, activated CD8^+^T cells, dendritic cell subtypes). Cell subtypes in some instances were aggregated (pneumocytes, bronchiolar epithelial cells, dendritic cell subtype). Activated CD8^+^ T cells were identified by the co-expression of CD8^+^ T cell markers and NK cell markers, along with too low read count to be considered doublets. To identify clusters consisting of cells undergoing rapid cell division, scMCA’s labeling of dividing cells was used. These clusters were combined into a single population named “Dividing Immune Cells”, as cells within this population expressed markers for both lymphocytes and myeloid cells. This cluster annotation was also corroborated with the CellCycleScoring() function in Seurat, where 97.7% of the cells in the dividing immune cell cluster were predicted to be in G2M (59.4%) or S (38.3%) phase. To account for invading cell types that were only present in MI samples, the interstitial macrophage and alveolar macrophage cells were removed from the cell type proportion calculations of the remaining cell types.

### Cell type-specific comparisons of MI to N and VI to N

DEGs were identified in scRNA-seq data for MI and VI comparisons to N in CD8^+^ T cells, NK cells, monocytes and DC subtypes, including pDCs and cDCs, using the non-parametric Wilcoxon rank sum test. The groups of DCα cells were incorporated into cDCs during DEG identification. Differential expression p-values were adjusted for multiple hypothesis testing using the Benjamini-Hochberg procedure (53). DEGs with FDR < 0.05 and absolute logFC > 0.1 for MI vs. N and VI vs. N comparisons were selected as significant. To study gene function and pathway enrichment among DEGs, golden hamster genes were mapped to homolog genes in the homo sapiens species using Biomart (http://www.ensembl.org/biomart). Significant genes were clustered into functional modules using Louvain community clustering based on the functional similarity between genes in the lung tissue, as predicted by HumanBase (https://hb.flatironinstitute.org) (53). Gene ontology (GO) term enrichment was performed for genes in each module. Significantly enriched terms with q-value < 0.05 were selected. Each module was annotated with distinct and representative GO terms. For CD8^+^ T cells, NK cells, and cDCs, gene functional clustering and enrichment analysis was performed for DEGs that were common to the MI and VI comparisons to N (Figure 5,C and D and Figure 6,C and D) and DEGs that are specific to either the MI or VI comparisons to N only (Supplemental Figure 10). For pDCs, however, functional clustering and enrichment analysis was only performed on MI vs. N given the insufficient number of DEGs in VI vs. N.

### Statistical analysis

Statistical analysis was performed using GraphPad Prism software, version 6. Two-way ANOVA with Tukey’s or Sidak’s corrections were respectively performed for multiple comparisons between vaccine groups or between time points. Significance between pre- and post- infection antibody titers was measured by multiple t tests with Holm Sidak’s correction for multiple comparisons (Supplemental Figure 4). ANOVA and post-hoc Tukey’s test pairwise comparisons were employed to determine the significance of scRNA-seq-based cell type proportion changes (Figure 3). Correlations were determined by two-sided Spearman s rank tests. \**P* < 0.05, \*\**P* < 0.01, \*\*\**P* < 0.001, \*\*\*\**P* < 0.0001.

### Data availability

RNA sequencing data have been deposited in NCBI’s Gene Expression Omnibus and are accessible through the GEO Series accession number GSE163838.

### CONTRIBUTIONS

M.M., D.E., A.C. and A.B, conceived and designed the study.

G.S-J. designed the vaccine construct.

C.E.M. performed the SARS-CoV-2 challenge experiment and animal procedures.

M.M. and C.P. processed hamster samples.

C.H., A.W. and L.M. performed ELISAs.

M.M. performed and analyzed SARS-CoV-2-based virology and serology assays.

P.R. performed single-cell RNA sequencing.

Y.W., G.R.S., A.B.R., X.C., W.C. analyzed single-cell RNA sequencing data.

K.W.B., M. Minai and B.M.N. performed histopathology and immunohistochemistry.

I.N.M. and S.P. analyzed the histopathology data.

I.N.M. analyzed the immunohistochemistry data.

P-Y.S. contributed a new reagent.

M.M., Y.W., G.R.S., A.B.R., X.C. and I.N.M. prepared manuscript figures and tables.

M.M., Y.W., G.R.S., A.B.R., X.C., W.C., I.N.M., I.R., O.G.T., E.Z., S.C.S. and A.B. wrote the manuscript.

D.E., E.Z., O.G.T., A.C., S.C.S. and A.B. supervised the research.

All authors reviewed, edited, and approved the final version of the manuscript.

## ACKNOWLEDGEMENTS

We thank members of the UTMB Animal Resource Center for technical assistance with the hamster experiment in ABSL-2 and ABSL-4 and husbandry support. We thank Steve Widen and the UTMB Next Generation Sequencing Core for quantification and pooling of scRNA-seq libraries for submission to the New York Genome Center. We thank the Anatomic Pathology Core facility at UTMB for embedding, sectioning and staining lung tissues for histopathology. The challenge virus used in this publication was generously provided by World Reference Center for Emerging Viruses and Arboviruses at UTMB. This work was supported by Moderna, Inc.

---

**Supplemental Figure 1.**
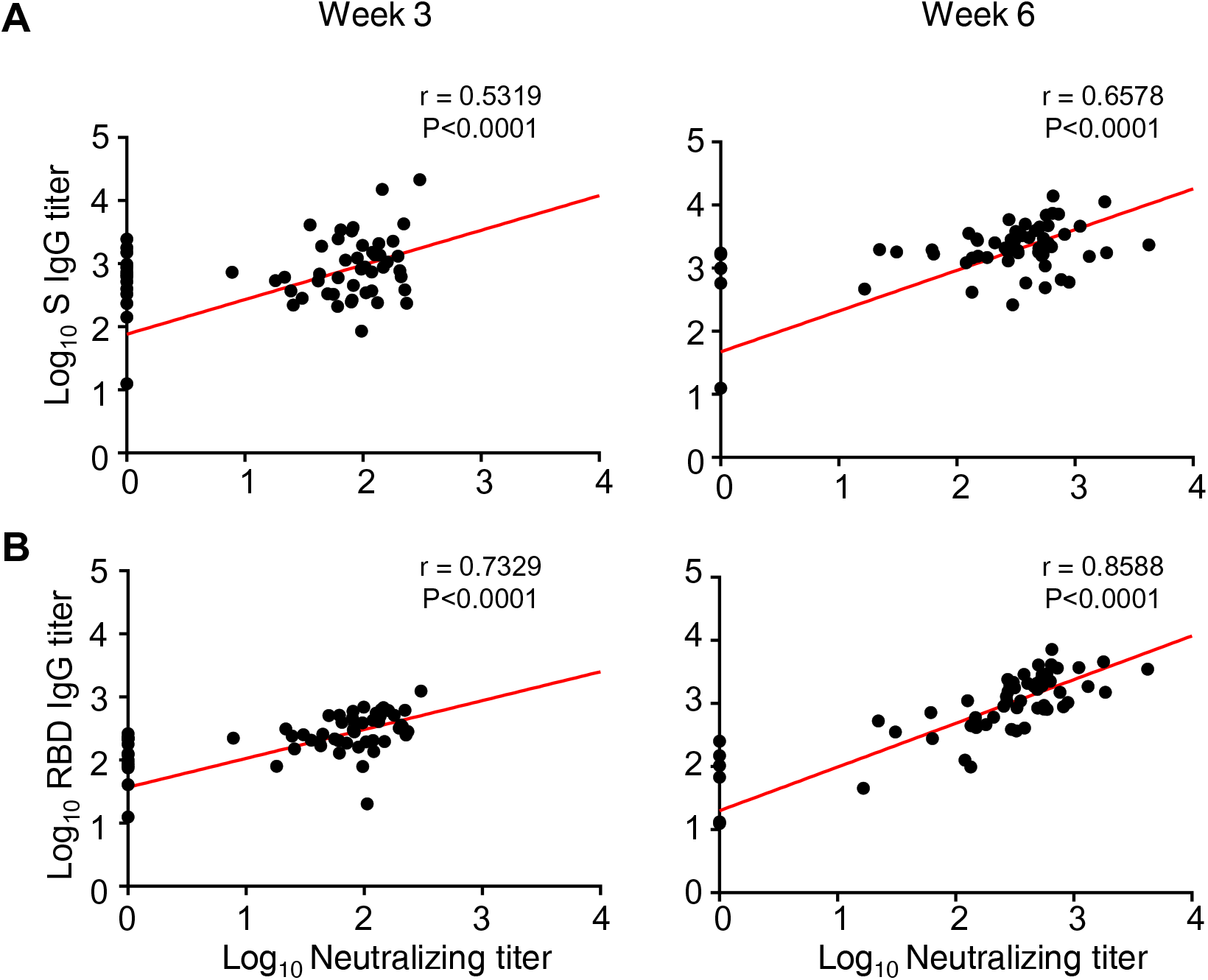
Correlation of antibody titers and neutralizing titers. **(A,B)** Correlations of S- (A) and RBD- (B) binding titers and neutralizing titers at weeks 3 and 6 post vaccination. Red line denote linear regression fit. P and **r** values determined by two-sided Spearman rank test.

**Supplemental Figure 2.**
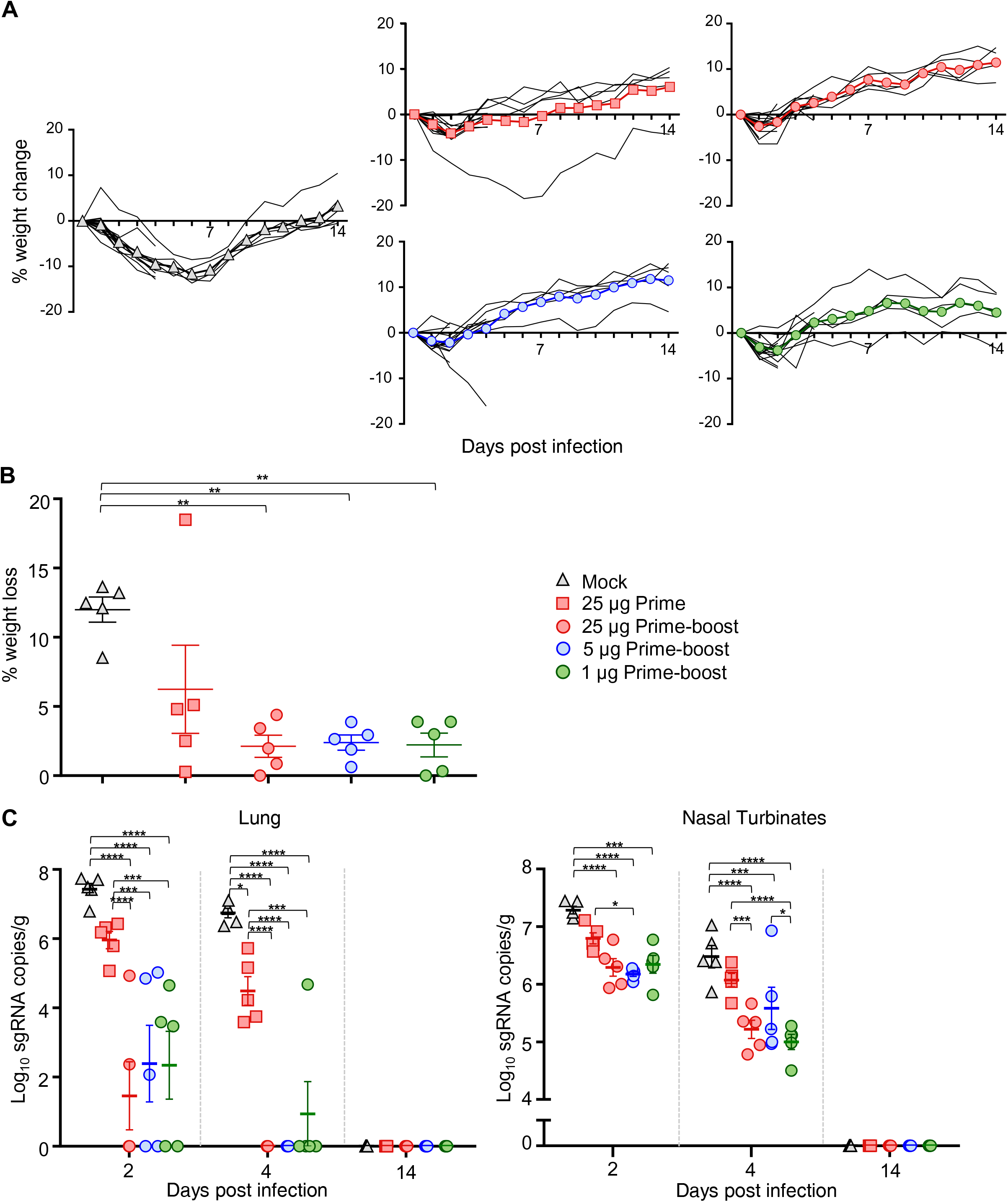
Characteristics of clinical disease following SARS-CoV-2 challenge of vaccinated hamsters. (A) Percent weight change in individual animals over 14 days post infection. Mean weight highlighted by line with color symbols. A weight measure for one hamster in the 25 ug prime-boost group at 2 dpi, the time of euthanasia, was excluded due to a measurement error. (B) Maximum weight loss in vaccinated groups, excluding animals serially euthanized on days 2 and 4 post challenge. (C) Viral sgRNA was measured by qRT-PCR in the lungs and nasal turbinates at serial endpoint days (2, 4 and 14 dpi). (B,C) Significance measured by ANOVA with Tukey’s correction for multiple comparisons (*P ≤ 0.05, **P ≤ 0.01, ***P ≤ 0.001, ****P ≤ 0.0001). Error bars represent ± SE.

**Supplemental Figure 3.**
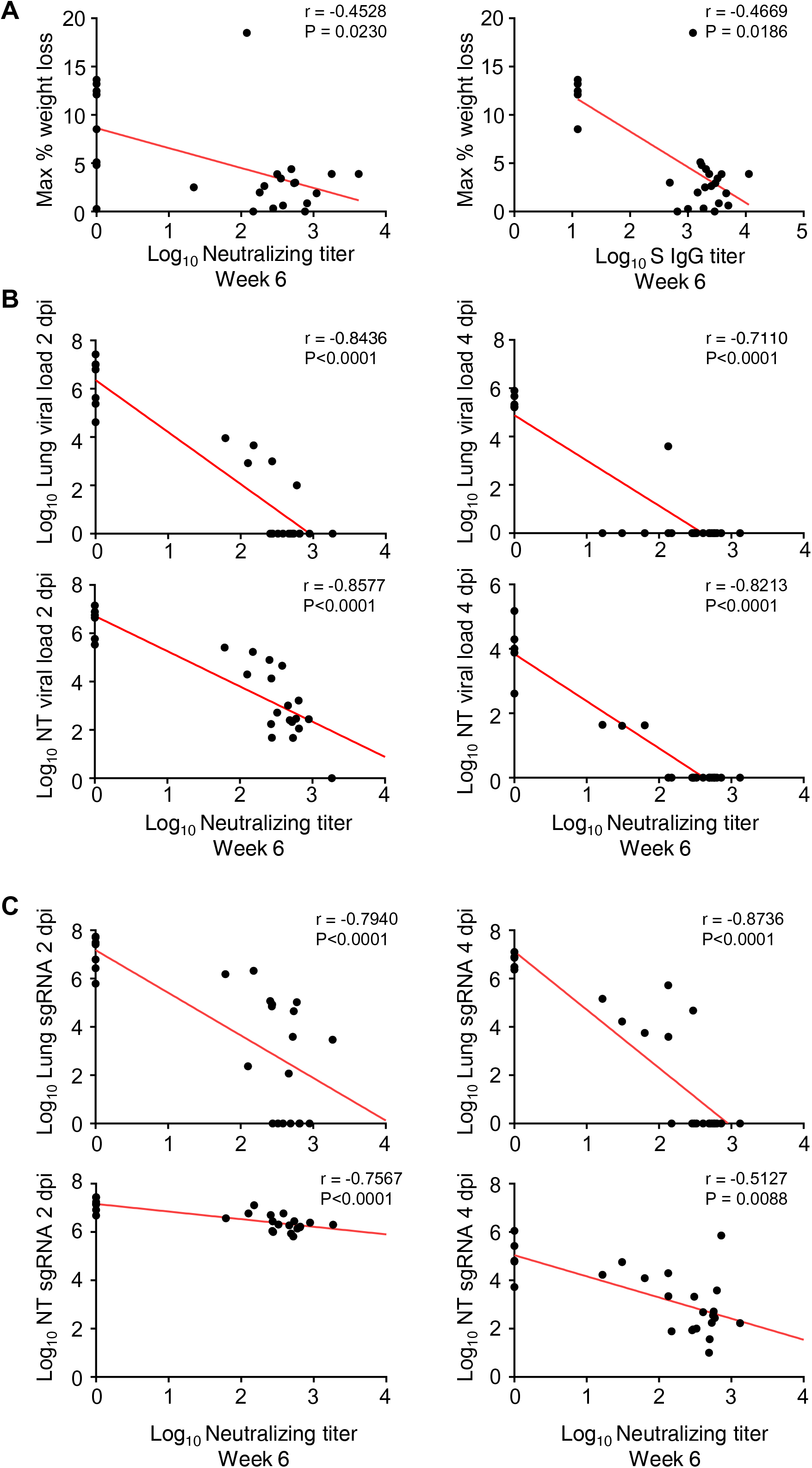
Correlation of antibody and neutralizing titers with disease parameters. (A) Correlations of S- binding titers and neutralizing titers at week 6 post vaccination with maximum percentage weight loss excluding hamsters euthanized at 2 and 4 dpi. Correlations between neutralizing titers at week 6 post vaccination and (B) virus load titers or (C) sgRNA in lungs and nasal turbinates at 2 and 4 dpi. Red line denote linear regression fit. P and **r** values determined by two-sided Spearman rank test

**Supplemental Figure 4.**
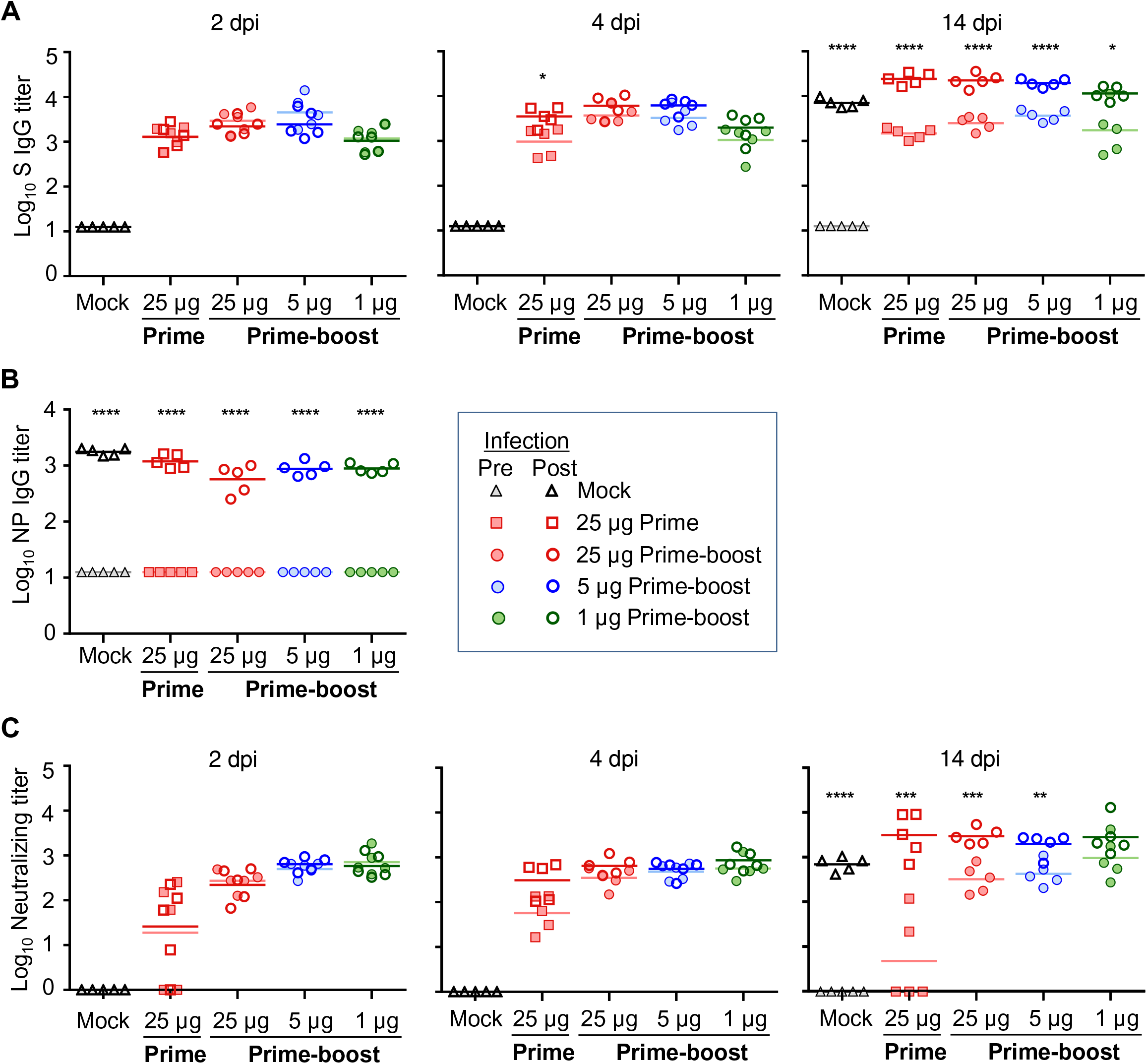
Serum antibody responses post SARS-CoV-2. challenge. (A) S-specific serum IgG titers in serum collected pre- (day 41 post-vaccination; shaded symbol) and post-infection (2, 4 and 14 days post-infection (dpi) open symbol), determined by ELISA. (A) S-specific serum IgG titers pre- (day 41 post-vaccination; shaded symbol) and post-infection (2, 4 and 14 days post-infection (dpi) open symbol) determined by ELISA. (B) NP-specific IgG titers in serum collected a day before challenge (day 41) and 14 dpi determined by ELISA. (C) Neutralizing titers in serum collected pre- (day 41 post-vaccination; shaded symbol) and post-infection (2, 4 and 14 days post-infection (dpi) open symbol), determined by plaque reduction assay. Bars denote group means. Significance between pre- and post- infection antibody titers measured by multiple t tests with Holm-Sidak’s correction for multiple comparisons (*P ≤ 0.05, **P ≤ 0.01, ***P ≤ 0.001, ****P ≤ 0.0001).

**Supplemental Figure 5.**
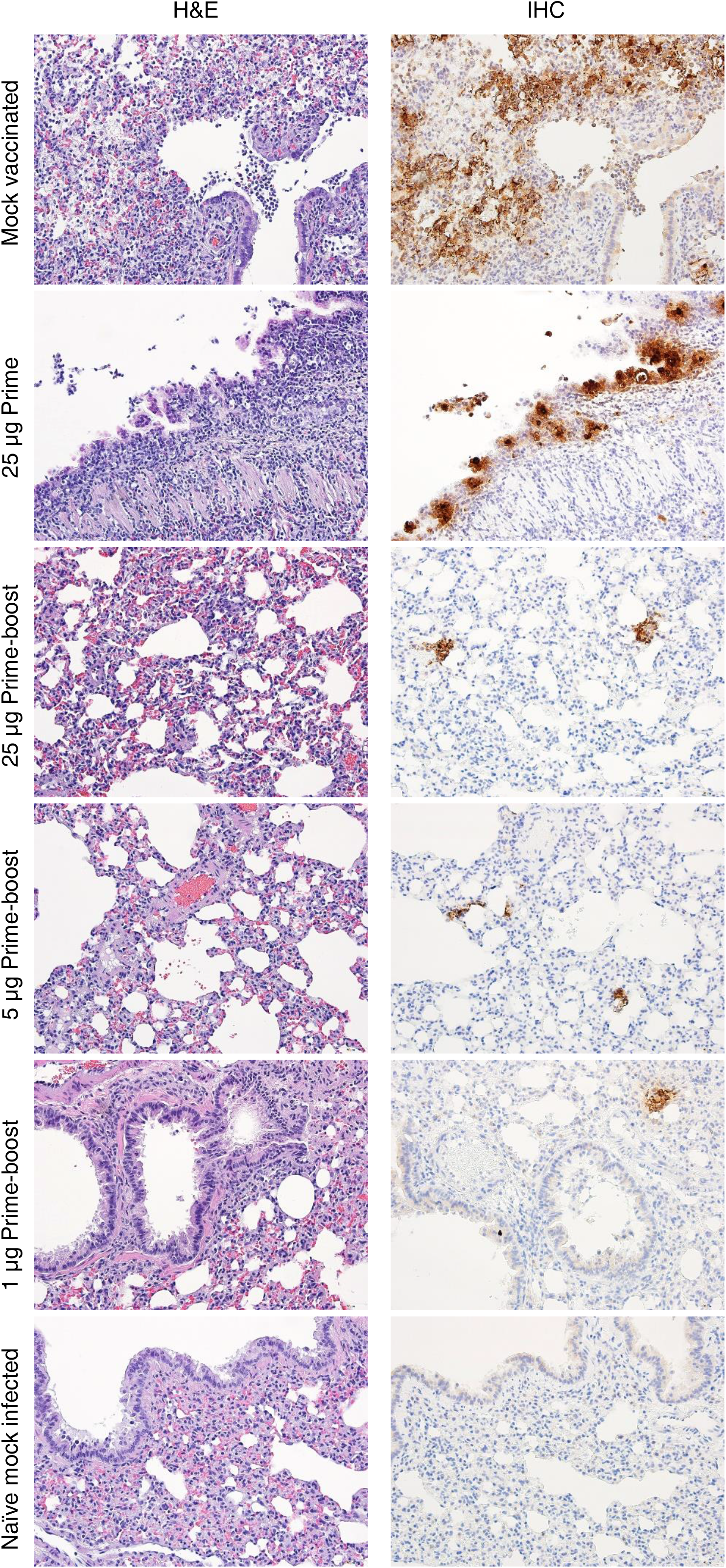
Disease following challenge of vaccinated hamsters with SARS-CoV-2 (continued from Fig. 2). Lung sections from mock vaccinated, prime-only and prime-boost vaccinated and naïve mock-infected groups at 4 dpi were stained with H&E, and representative photomicrographs (original magnification ×20 (scale bars, 50 µm) as indicated) from each group with virus antigen (arrowhead) in lungs, stained by IHC.

**Supplemental Figure 6.**
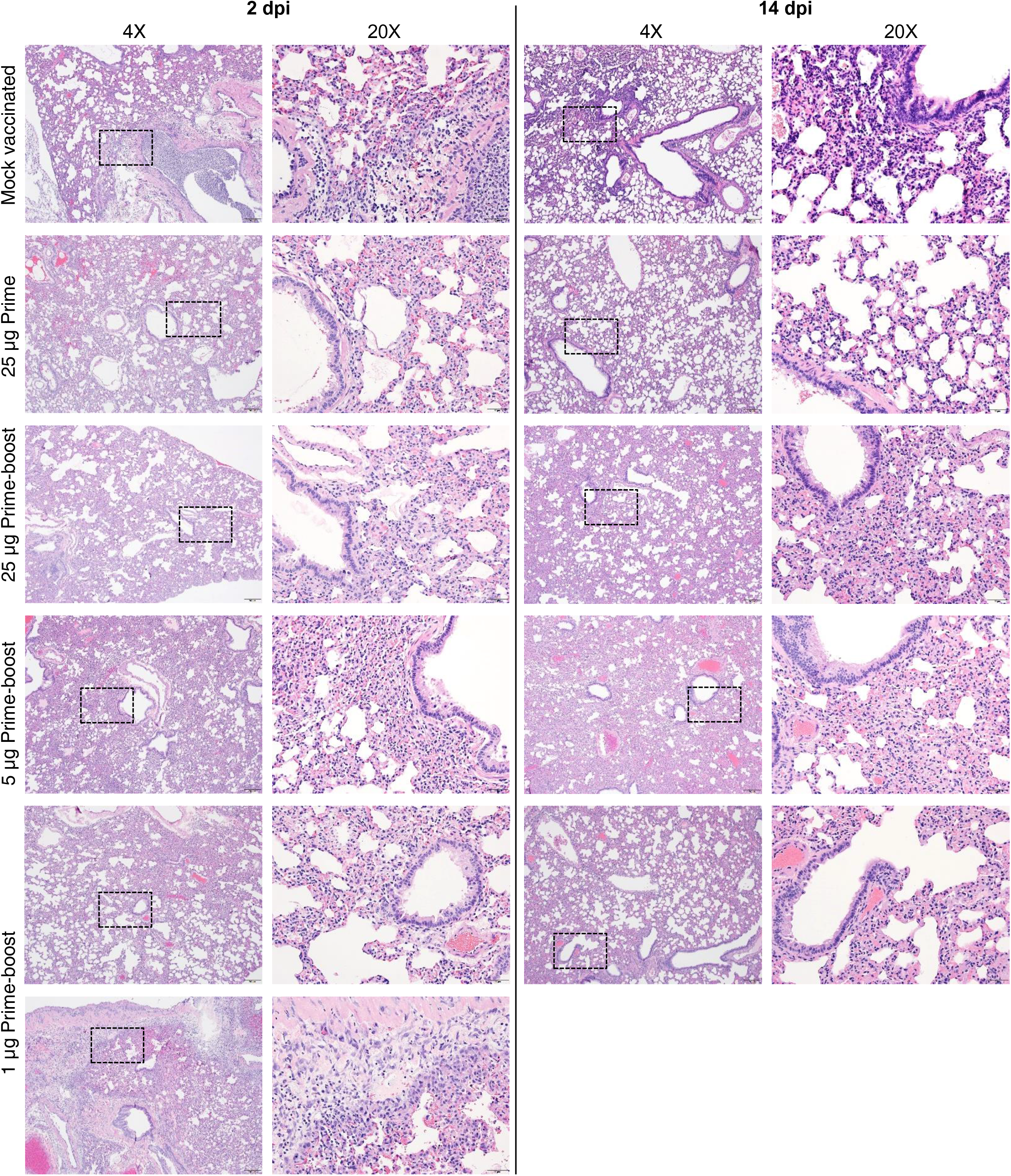
Disease following challenge of vaccinated hamsters with SARS-CoV-2 (continued from Fig 2). Lung sections from mock vaccinated, prime-only and prime-boost vaccinated animals at 2 and 14 dpi were stained with H&E, and representative photomicrographs (original magnification ×4 (scale bars, 200 µm) and ×20 (scale bars, 50 µm) as indicated) from each group with virus antigen (arrowhead) in lungs, stained by IHC, are shown.

**Supplemental Figure 7.**
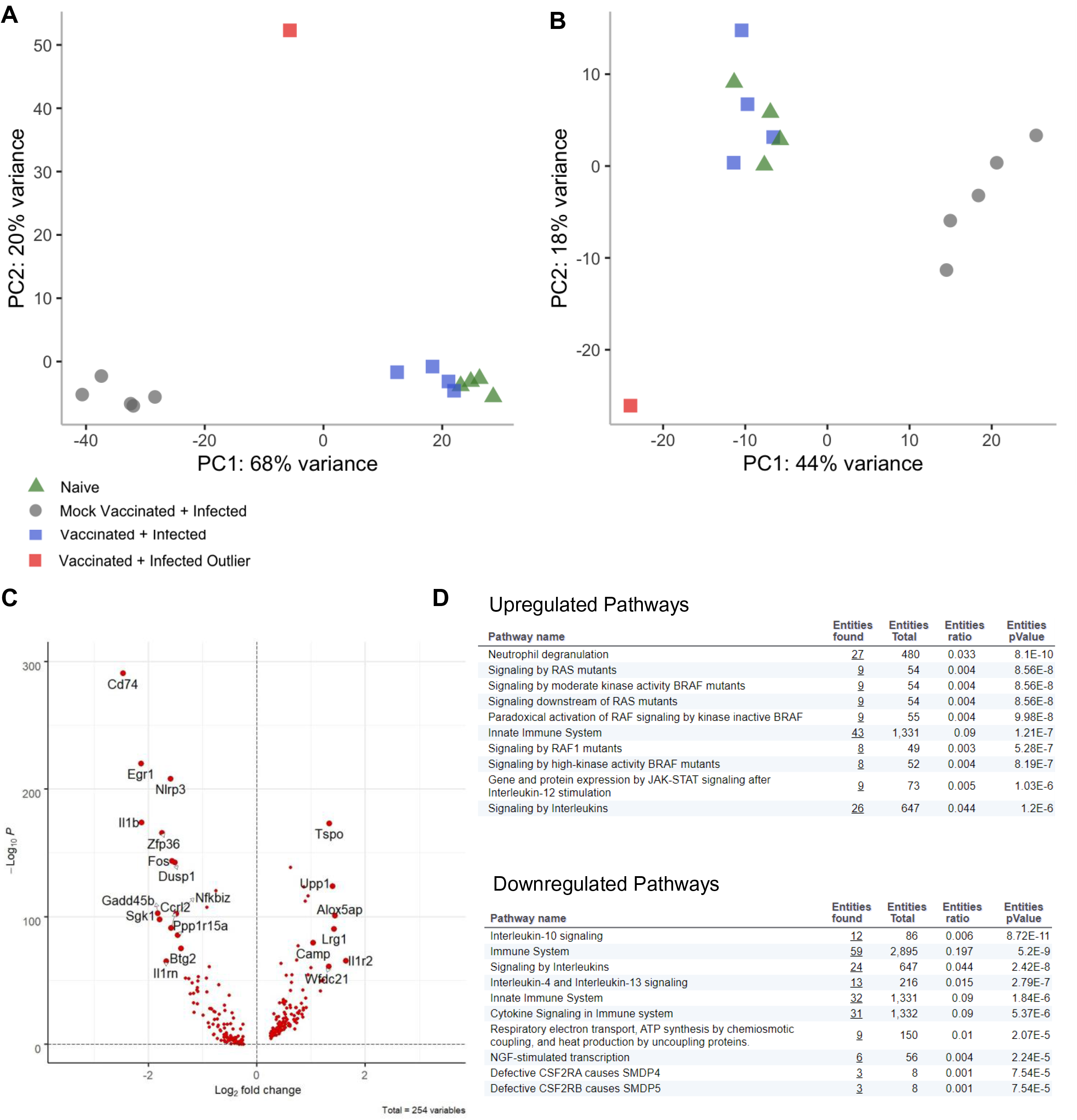
Gene regulation in the vaccinated infected outlier sample. (A,B) PCA of pseudo-bulk RNA-seq data including all cell types (A) and just granulocytes (B) Colors indicate groups. Outlier shown in red. (C) Volcano plot of genes that are differentially expressed (DE) between the outlier sample and the other VI samples. 270 differentially expressed genes satisfying a logFC threshold of 0.25 and an FDR adjusted p-value of 0.05 were identified: 158 up-regulated, 112 down-regulated. Only DE genes are shown. (D) Upregulated and downregulated Reactome pathways for the outlier vs. the other VI samples.

**Supplemental Figure 8.**
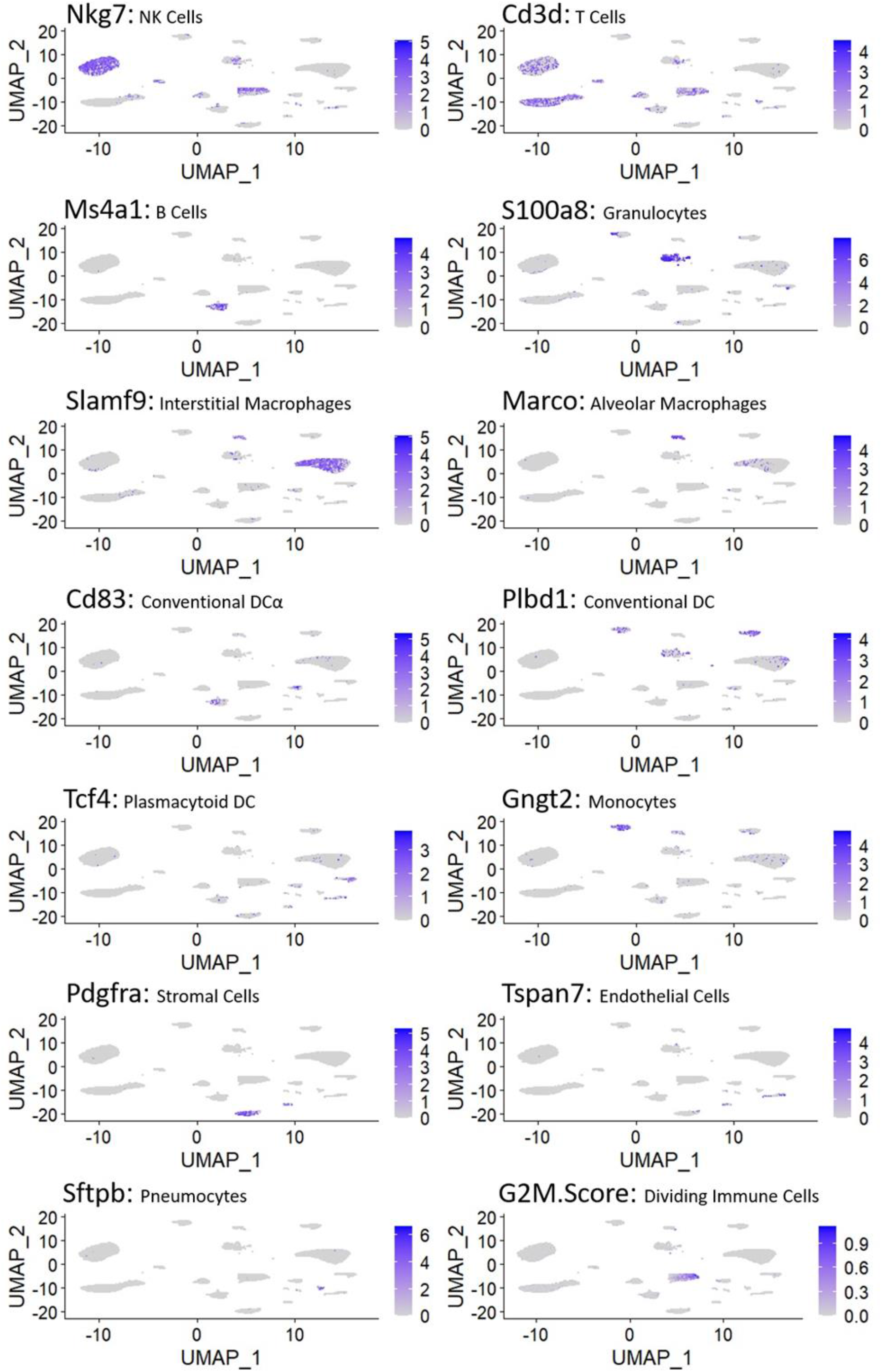
Cell type-specific markers differentiate single cell clusters. Each plot displays the expression pattern of a representative cell type-specific marker on the integrated UMAP plot. Feature plots are labeled by cell-type marker and corresponding cell type. Dividing immune cells are marked by the G2M score derived in Seurat.

**Supplemental Figure 9.**
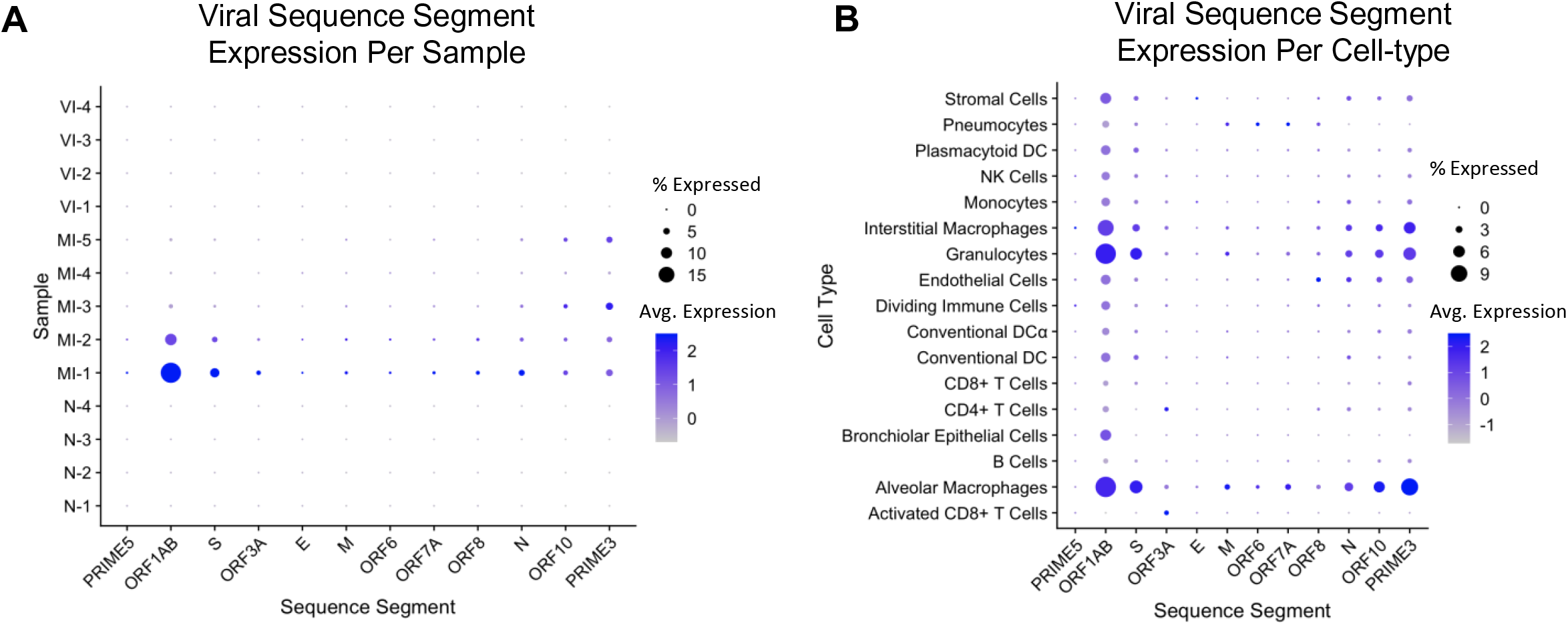
Expression of SARS-CoV-2 genes. (A) Dot plot showing expression of each viral sequence segment in each sample. PRIME5 and PRIME3 represent the 5’ and 3’ ends, respectively, of the viral genome. (B) Dot plot showing expression of each viral sequence segment in each cell type (including MI cells only).

**Supplemental Figure 10.**
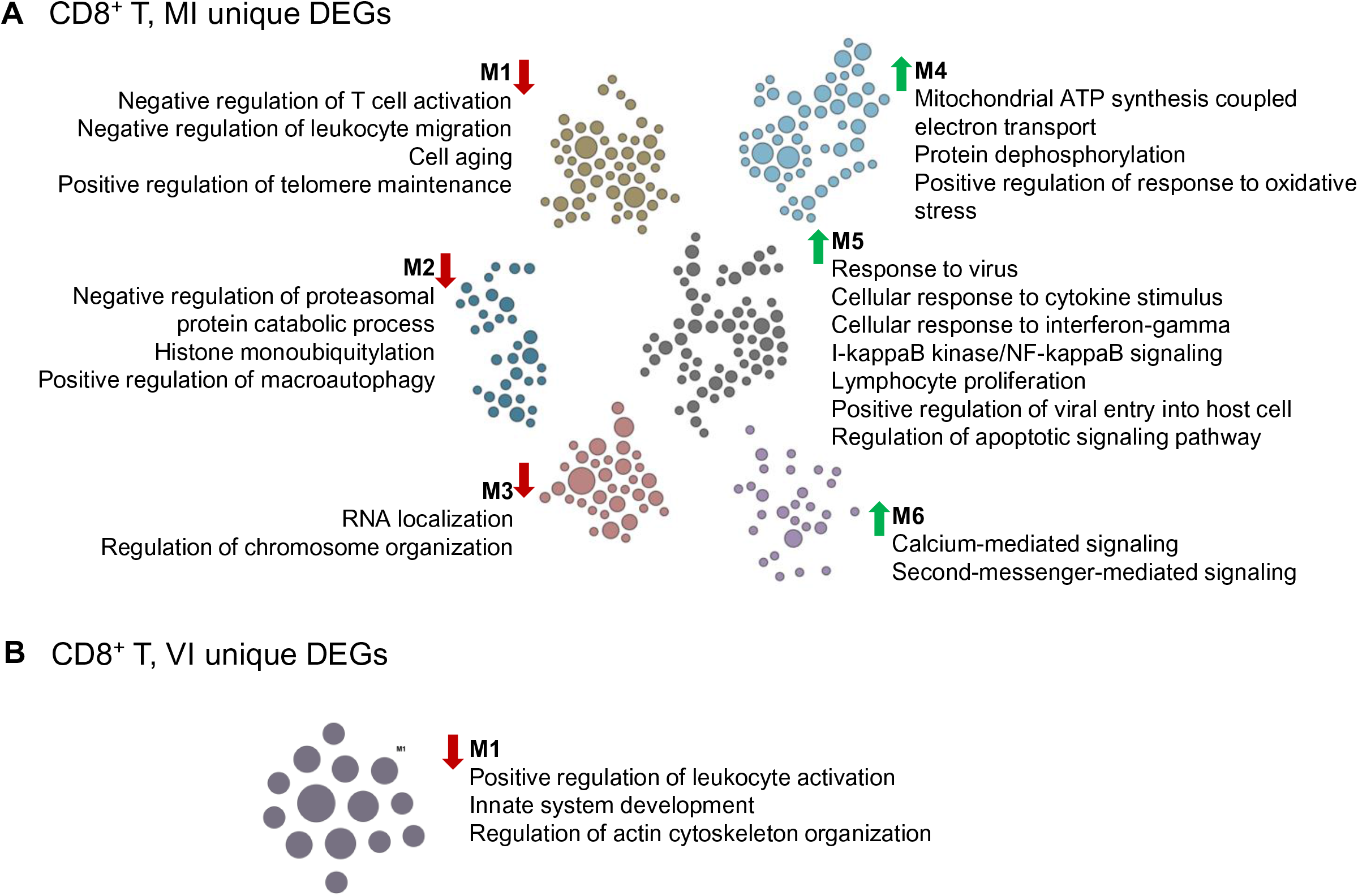
Functional pathway analysis of unique MI vs. N and VI vs. N comparisons in CD8^+^ T cells. (A,B) Functional network showing modules of enriched pathways using DEGs specific to the (A) MI vs. N or (B) VI vs. N comparison in CD8^+^ T cells, i.e. fall outside the Venn diagram overlap in Fig. 5A. Each gene is represented by a circle and color coded according to module. The size of each circle reflects its connectivity in the network. Edges are not shown, allowing for easy viewing. The module label is shown with the functional processes and pathways identified in each module. Up or down regulation of pathway is indicated by arrow direction.

**Supplemental Figure 11.**
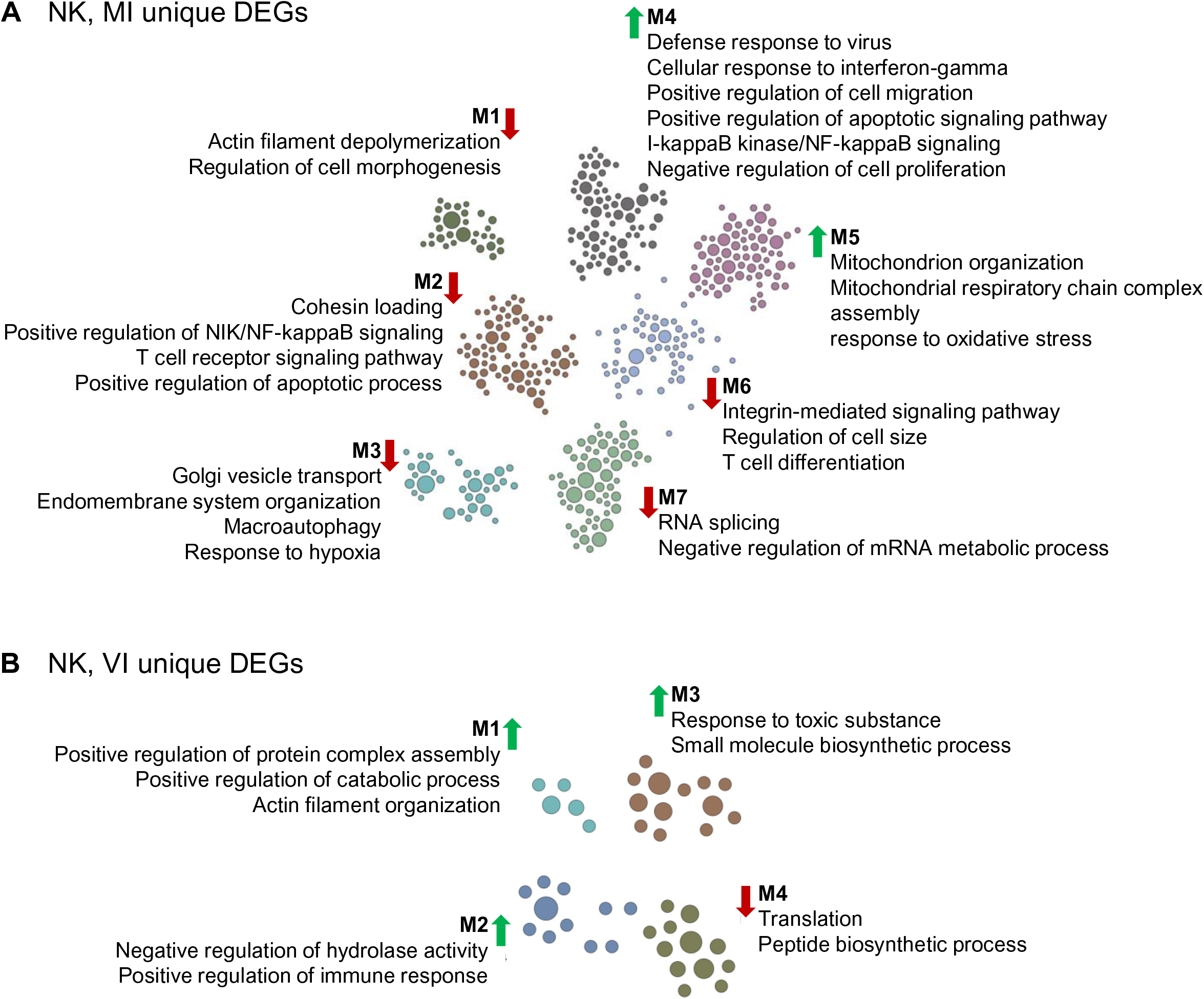
Functional pathway analysis of unique MI vs. N and VI vs. N comparisons in NK cells. (A,B) Functional network showing modules of enriched pathways using DEGs specific to the (A) MI vs. N or (B) VI vs. N comparison in NK cells, i.e. fall outside the Venn diagram overlap in Fig. 5B. Each gene is represented by a circle and color coded according to module. The size of each circle reflects its connectivity in the network. Edges are not shown, allowing for easy viewing. The module label is shown with the functional processes and pathways identified in each module. Up or down regulation of pathway is indicated by arrow direction.

**Supplemental Figure 12.**
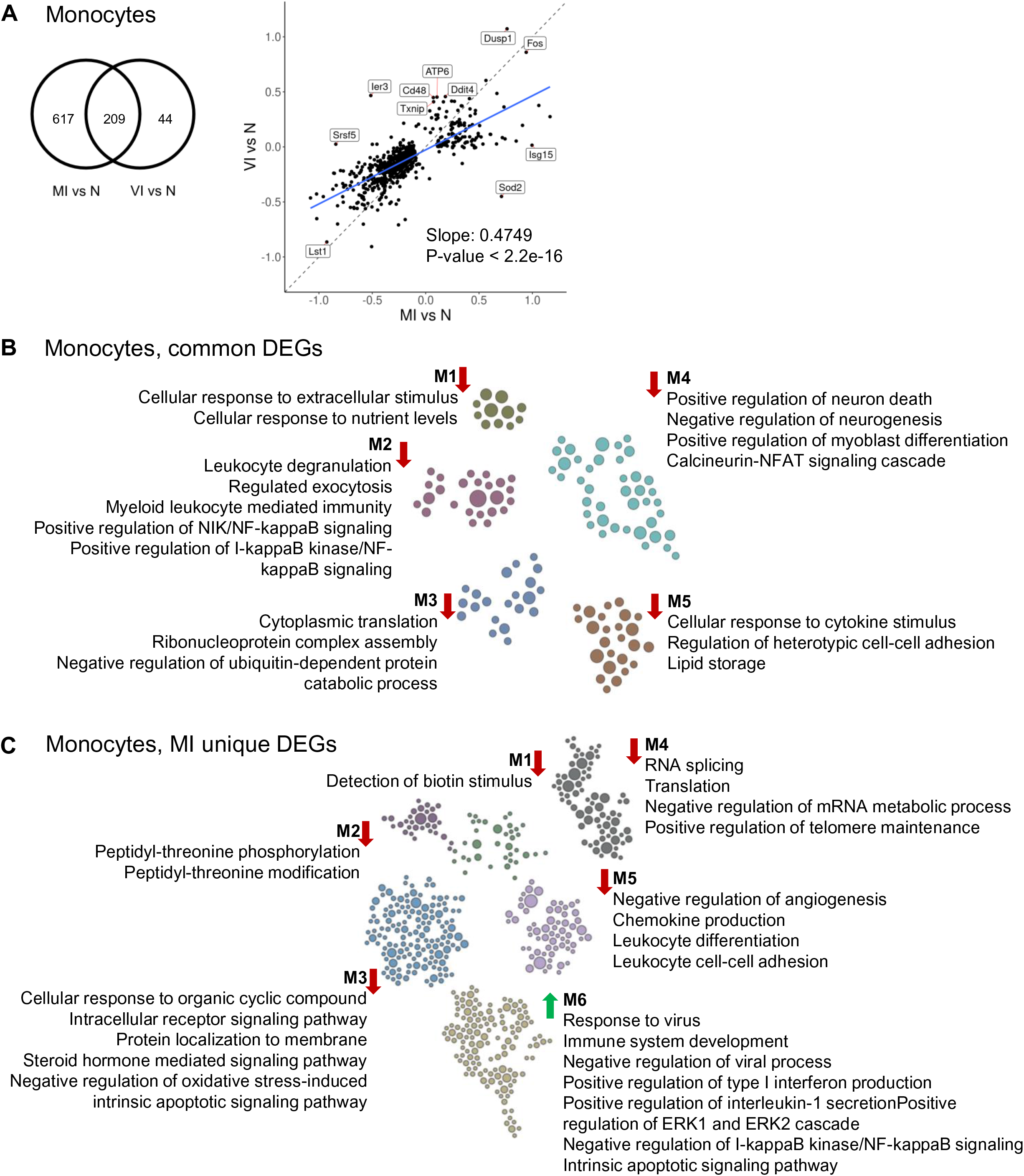
Functional pathway analysis of MI vs. N and VI vs. N comparisons in monocytes. (A) Venn diagram of the union of DEGs of the two comparisons in monocytes. Scatter plot of the logFC of the union of DEGs of the two comparisons in Monocytes, i.e. the same group of genes as in the Venn diagram. For each comparison, DEGs were selected with thresholds FDR < 0.05 and absolute logFC > 0.1. Linear regression model was fitted to the scatter plot (adjusted R^∧^2 = 0.5779). (B,C) Functional network showing modules of enriched pathways using (B) DEGs shared between two comparisons in monocytes or (C) DEGs unique to MI monocytes. Each gene is represented by a circle and color coded according to module. The size of each circle reflects its connectivity in the network. Edges are not shown, allowing for easy viewing. The module label is shown with the functional processes and pathways identified in each module. Up or down regulation of pathway is indicated by arrow direction.

**Supplemental Figure 13.**
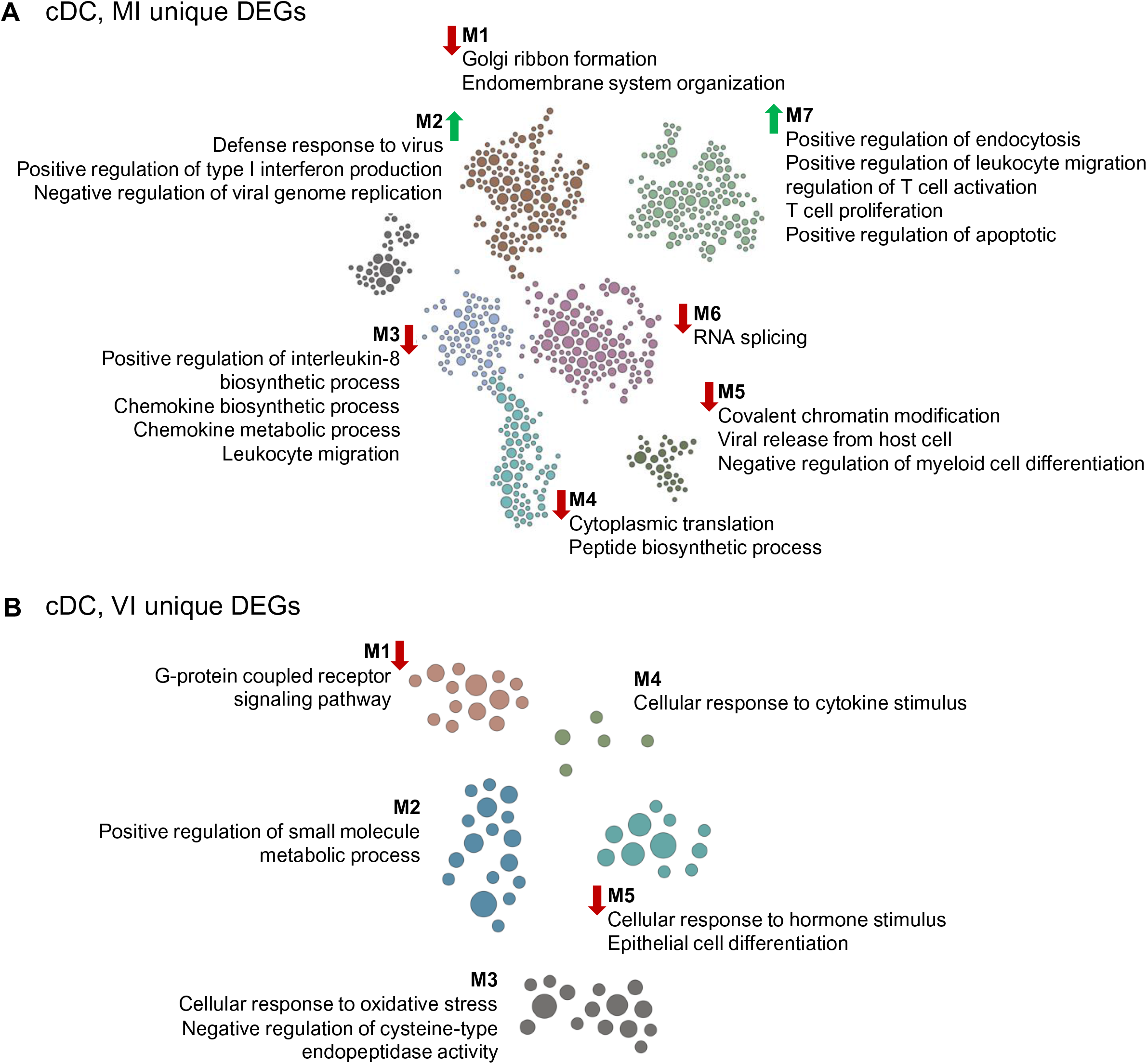
Functional pathway analysis of unique MI vs. N and VI vs. N comparisons in cDC. (A,B) Functional network showing modules of enriched pathways using DEGs specific to the (A) MI vs. N or (B) VI vs. N comparison in cDC, i.e. fall outside the Venn diagram overlap in Fig. 6A. Each gene is represented by a circle and color coded according to module. The size of each circle reflects its connectivity in the network. Edges are not shown, allowing for easy viewing. The module label is shown with the functional processes and pathways identified in each module. Up or down regulation of pathway is indicated by arrow direction.

**Supplemental Table 1.**
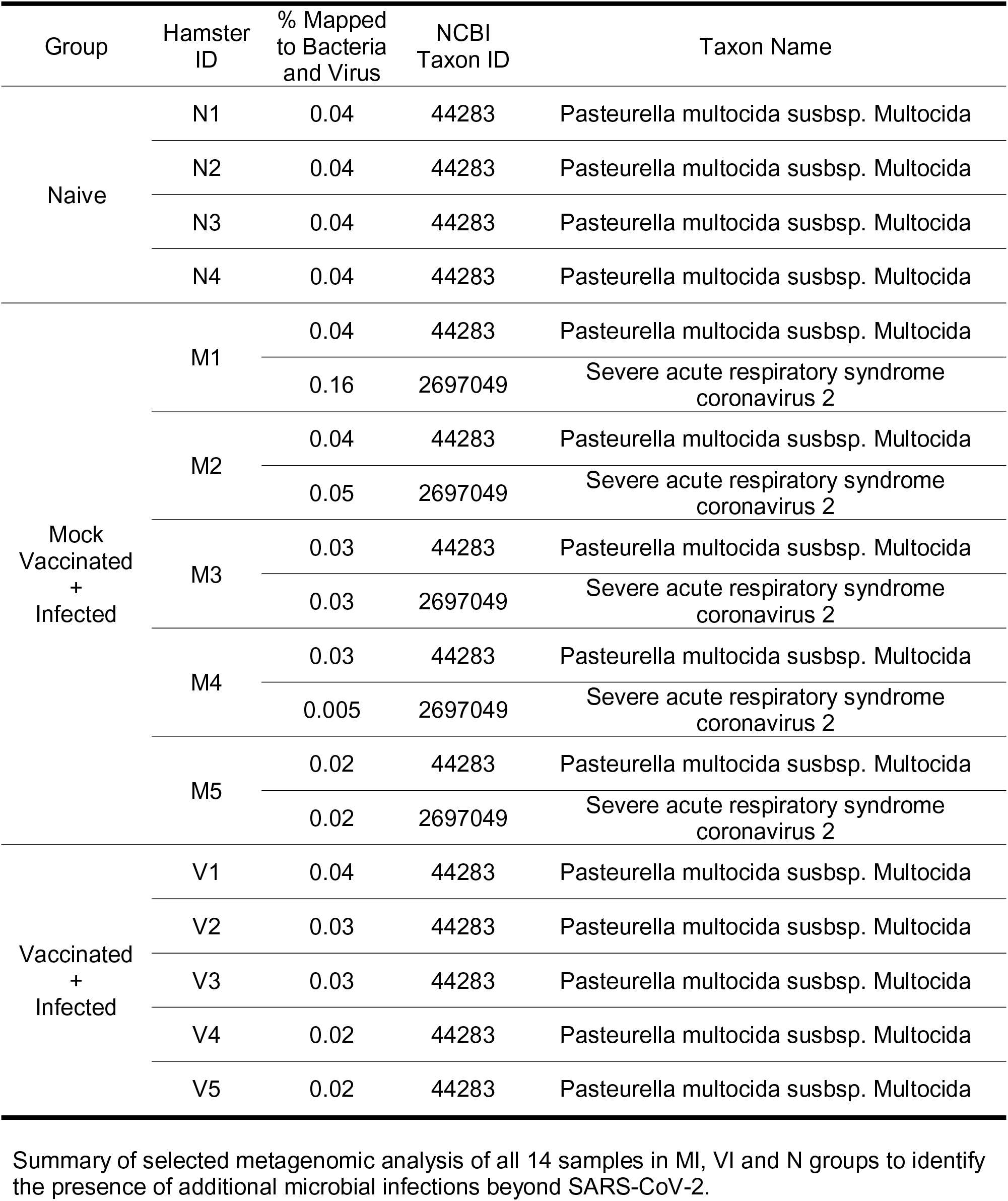
Metagenomic Analysis of all 14 samples: selected classification results

| Group | Hamster ID | % Mapped to Bacteria and Virus | NCBI Taxon ID | Taxon Name |
| --- | --- | --- | --- | --- |
| Naive | N1 | 0.04 | 44283 | Pasteurella multocida subsp. Multocida |
|  | N2 | 0.04 | 44283 | Pasteurella multocida subsp. Multocida |
|  | N3 | 0.04 | 44283 | Pasteurella multocida subsp. Multocida |
|  | N4 | 0.04 | 44283 | Pasteurella multocida subsp. Multocida |
| Mock Vaccinated + Infected | M1 | 0.04 | 44283 | Pasteurella multocida subsp. Multocida |
|  |  | 0.16 | 2697049 | Severe acute respiratory syndrome coronavirus 2 |
|  | M2 | 0.04 | 44283 | Pasteurella multocida subsp. Multocida |
|  |  | 0.05 | 2697049 | Severe acute respiratory syndrome coronavirus 2 |
|  | M3 | 0.03 | 44283 | Pasteurella multocida subsp. Multocida |
|  |  | 0.03 | 2697049 | Severe acute respiratory syndrome coronavirus 2 |
|  | M4 | 0.03 | 44283 | Pasteurella multocida subsp. Multocida |
|  |  | 0.005 | 2697049 | Severe acute respiratory syndrome coronavirus 2 |
|  | M5 | 0.02 | 44283 | Pasteurella multocida subsp. Multocida |
|  |  | 0.02 | 2697049 | Severe acute respiratory syndrome coronavirus 2 |
| Vaccinated + Infected | V1 | 0.04 | 44283 | Pasteurella multocida subsp. Multocida |
|  | V2 | 0.03 | 44283 | Pasteurella multocida subsp. Multocida |
|  | V3 | 0.03 | 44283 | Pasteurella multocida subsp. Multocida |
|  | V4 | 0.02 | 44283 | Pasteurella multocida subsp. Multocida |
|  | V5 | 0.02 | 44283 | Pasteurella multocida subsp. Multocida |

Summary of selected metagenomic analysis of all 14 samples in MI, VI and N groups to identify the presence of additional microbial infections beyond SARS-CoV-2.

**Supplemental Table 2.**
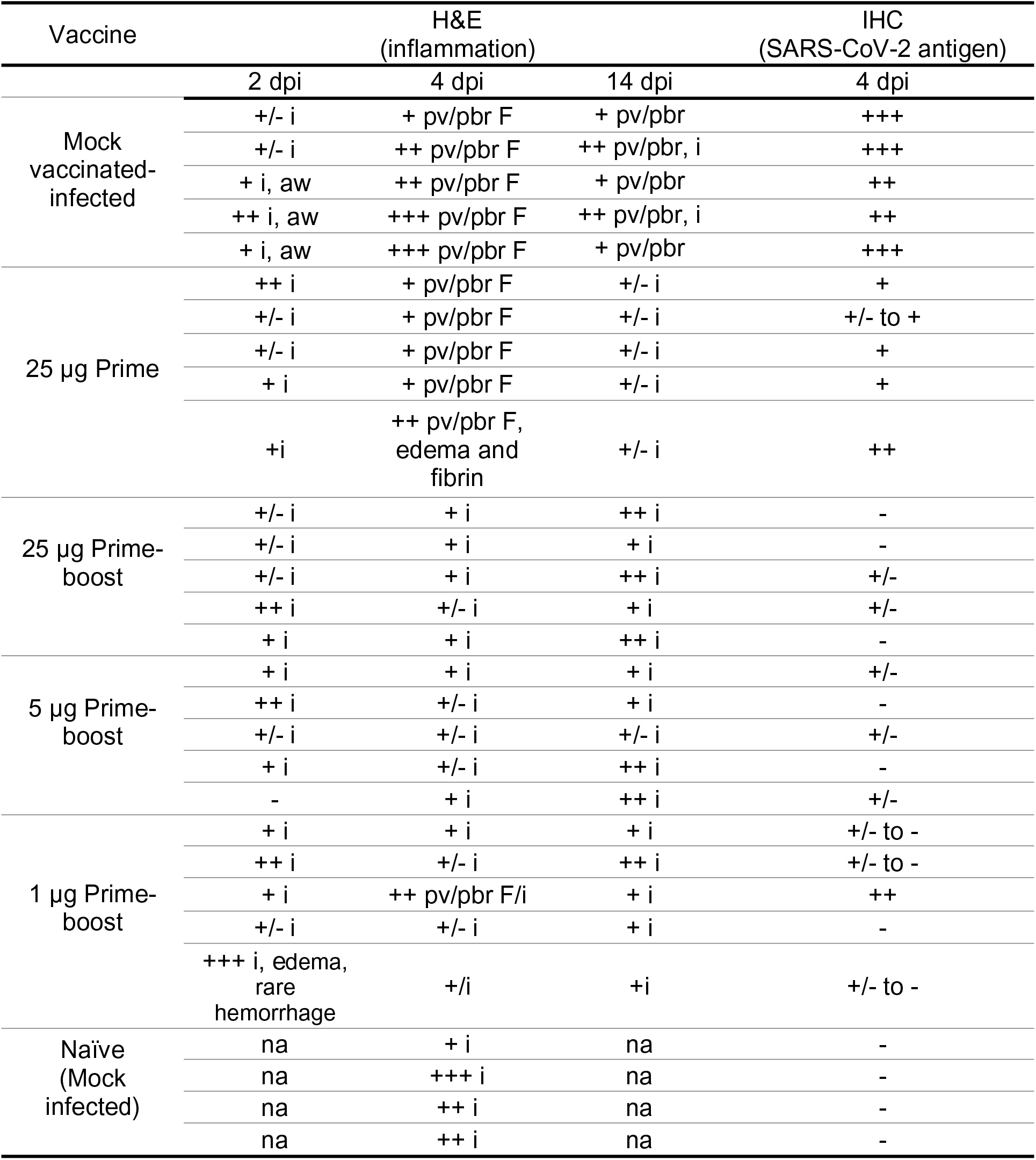
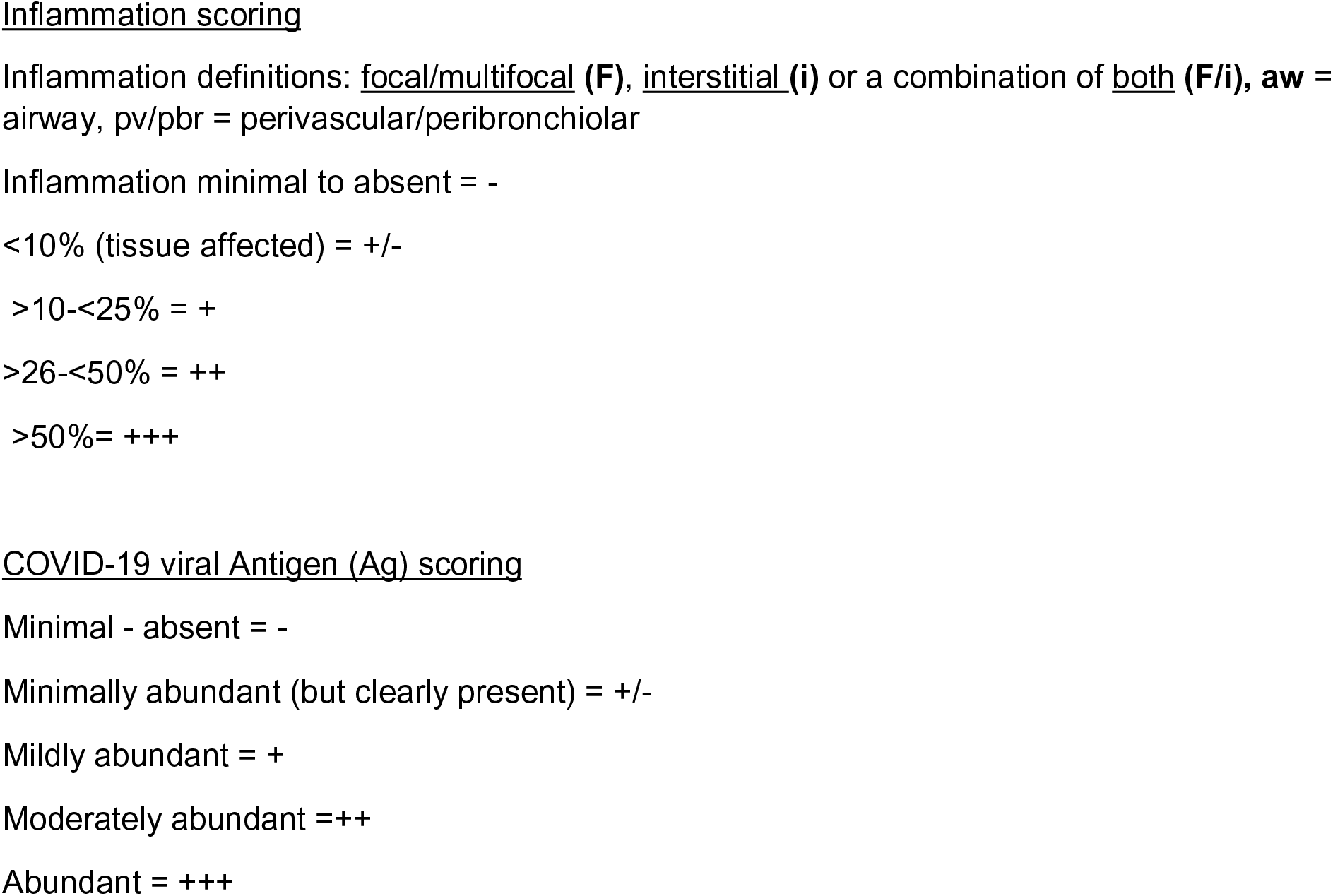
Inflammation and viral antigen in lungs of mRNA-1273 vaccinated hamsters following SARS-CoV-2 infection

## REFERENCES

1. Corbett KS, Edwards DK, Leist SR, Abiona OM, Boyoglu-Barnum S, Gillespie RA, Himansu S, Schafer A, Ziwawo CT, DiPiazza AT, et al. SARS-CoV-2 mRNA vaccine design enabled by prototype pathogen preparedness. Nature. 2020.

2. Meyer M, Huang E, Yuzhakov O, Ramanathan P, Ciaramella G, and Bukreyev A. Modified mRNA-Based Vaccines Elicit Robust Immune Responses and Protect Guinea Pigs From Ebola Virus Disease. J Infect Dis. 2018;217(3):451–5.

3. Feldman RA, Fuhr R, Smolenov I, Mick Ribeiro A, Panther L, Watson M, Senn JJ, Smith M, Almarsson, Pujar HS, et al. mRNA vaccines against H10N8 and H7N9 influenza viruses of pandemic potential are immunogenic and well tolerated in healthy adults in phase 1 randomized clinical trials. Vaccine. 2019;37(25):3326–34.

4. Aliprantis AO, Shaw CA, Griffin P, Farinola N, Railkar RA, Cao X, Liu W, Sachs JR, Swenson CJ, Lee H, et al. A phase 1, randomized, placebo-controlled study to evaluate the safety and immunogenicity of an mRNA-based RSV prefusion F protein vaccine in healthy younger and older adults. Hum Vaccin Immunother. 2020:1–14.

5. ClinicalTrials.gov. A Phase 1, Randomized, Observer-Blind, Placebo-Controlled, Dose- Ranging Study to Evaluate the Safety, Reactogenicity, and Immunogenicity of Cytomegalovirus Vaccines mRNA-1647 and mRNA-1443 When Administered to Healthy Adults. https://clinicaltrials.gov/ct2/show/NCT03382405.

6. ClinicalTrials.gov. A Phase 1, Randomized, Observer-Blind, Placebo-Controlled, Dose- Ranging Study to Evaluate the Safety, Reactogenicity, and Immunogenicity of mRNA- 1653, a Combined Human Metapneumovirus and Human Parainfluenza Virus Type 3 Vaccine, When Administered to Healthy Adults. https://clinicaltrials.gov/ct2/show/NCT03392389.

7. ClinicalTrials.gov. A Phase 1, Randomized, Observer-Blind, Placebo-Controlled, Dose- Ranging Study to Evaluate the Safety, Tolerability, and Immunogenicity of Zika Vaccine mRNA-1893 in Healthy Flavivirus Seropositive and Seronegative Adults. https://clinicaltrials.gov/ct2/show/NCT04064905.

8. Corbett KS, Flynn B, Foulds KE, Francica JR, Boyoglu-Barnum S, Werner AP, Flach B, O’Connell S, Bock KW, Minai M, et al. Evaluation of the mRNA-1273 Vaccine against SARS-CoV-2 in Nonhuman Primates. N Engl J Med. 2020.

9. Jackson LA, Anderson EJ, Rouphael NG, Roberts PC, Makhene M, Coler RN, McCullough MP, Chappell JD, Denison MR, Stevens LJ, et al. An mRNA Vaccine against SARS-CoV-2 - Preliminary Report. N Engl J Med. 2020.

10. Anderson EJ, Rouphael NG, Widge AT, Jackson LA, Roberts PC, Makhene M, Chappell JD, Denison MR, Stevens LJ, Pruijssers AJ, et al. Safety and Immunogenicity of SARS- CoV-2 mRNA-1273 Vaccine in Older Adults. N Engl J Med. 2020.

11. Baden LR, El Sahly HM, Essink B, Kotloff K, Frey S, Novak R, Diemert D, Spector SA, Rouphael N, Creech CB, et al. Efficacy and Safety of the mRNA-1273 SARS-CoV-2 Vaccine. N Engl J Med. 2020.

12. Munster VJ, Feldmann F, Williamson BN, van Doremalen N, Perez-Perez L, Schulz J, Meade-White K, Okumura A, Callison J, Brumbaugh B, et al. Respiratory disease in rhesus macaques inoculated with SARS-CoV-2. Nature. 2020;585(7824):268–72.

13. Woolsey C, Borisevich V, Prasad AN, Agans KN, Deer DJ, Dobias NS, Heymann JC, Foster SL, Levine CB, Medina L, et al. Establishment of an African green monkey model for COVID-19. bioRxiv. 2020.

14. Rockx B, Kuiken T, Herfst S, Bestebroer T, Lamers MM, Oude Munnink BB, de Meulder D, van Amerongen G, van den Brand J, Okba NMA, et al. Comparative pathogenesis of COVID-19, MERS, and SARS in a nonhuman primate model. Science. 2020;368(6494):1012–5.

15. Shi J, Wen Z, Zhong G, Yang H, Wang C, Huang B, Liu R, He X, Shuai L, Sun Z, et al. Susceptibility of ferrets, cats, dogs, and other domesticated animals to SARS- coronavirus 2. Science. 2020;368(6494):1016–20.

16. Kim YI, Kim SG, Kim SM, Kim EH, Park SJ, Yu KM, Chang JH, Kim EJ, Lee S, Casel MAB, et al. Infection and Rapid Transmission of SARS-CoV-2 in Ferrets. Cell Host Microbe. 2020;27(5):704–9 e2.

17. Blanco-Melo D, Nilsson-Payant BE, Liu WC, Uhl S, Hoagland D, Moller R, Jordan TX, Oishi K, Panis M, Sachs D, et al. Imbalanced Host Response to SARS-CoV-2 Drives Development of COVID-19. Cell. 2020;181(5):1036–45 e9.

18. Dinnon KH, 3rd, Leist SR, Schafer A, Edwards CE, Martinez DR, Montgomery SA, West A, Yount BL, Jr., Hou YJ, Adams LE, et al. A mouse-adapted model of SARS-CoV-2 to test COVID-19 countermeasures. Nature. 2020.

19. Imai M, Iwatsuki-Horimoto K, Hatta M, Loeber S, Halfmann PJ, Nakajima N, Watanabe T, Ujie M, Takahashi K, Ito M, et al. Syrian hamsters as a small animal model for SARS- CoV-2 infection and countermeasure development. Proc Natl Acad Sci U S A. 2020;117(28):16587–95.

20. Chan JF, Zhang AJ, Yuan S, Poon VK, Chan CC, Lee AC, Chan WM, Fan Z, Tsoi HW, Wen L, et al. Simulation of the clinical and pathological manifestations of Coronavirus Disease 2019 (COVID-19) in golden Syrian hamster model: implications for disease pathogenesis and transmissibility. Clin Infect Dis. 2020.

21. Tostanoski LH, Wegmann F, Martinot AJ, Loos C, McMahan K, Mercado NB, Yu J, Chan CN, Bondoc S, Starke CE, et al. Ad26 vaccine protects against SARS-CoV-2 severe clinical disease in hamsters. Nat Med. 2020.

22. Sia SF, Yan LM, Chin AWH, Fung K, Choy KT, Wong AYL, Kaewpreedee P, Perera R, Poon LLM, Nicholls JM, et al. Pathogenesis and transmission of SARS-CoV-2 in golden hamsters. Nature. 2020;583(7818):834–8.

23. Winkler ES, Bailey AL, Kafai NM, Nair S, McCune BT, Yu J, Fox JM, Chen RE, Earnest JT, Keeler SP, et al. SARS-CoV-2 infection of human ACE2-transgenic mice causes severe lung inflammation and impaired function. Nat Immunol. 2020.

24. Butler A, Hoffman P, Smibert P, Papalexi E, and Satija R. Integrating single-cell transcriptomic data across different conditions, technologies, and species. Nat Biotechnol. 2018;36(5):411–20.

25. Liao M, Liu Y, Yuan J, Wen Y, Xu G, Zhao J, Cheng L, Li J, Wang X, Wang F, et al. Single-cell landscape of bronchoalveolar immune cells in patients with COVID-19. Nat Med. 2020;26(6):842–4.

26. Chua RL, Lukassen S, Trump S, Hennig BP, Wendisch D, Pott F, Debnath O, Thurmann L, Kurth F, Volker MT, et al. COVID-19 severity correlates with airway epithelium-immune cell interactions identified by single-cell analysis. Nat Biotechnol. 2020;38(8):970–9.

27. Zhang JY, Wang XM, Xing X, Xu Z, Zhang C, Song JW, Fan X, Xia P, Fu JL, Wang SY, et al. Single-cell landscape of immunological responses in patients with COVID-19. Nat Immunol. 2020;21(9):1107–18.

28. Berlin DA, Gulick RM, and Martinez FJ. Severe Covid-19. N Engl J Med. 2020.

29. Maucourant C, Filipovic I, Ponzetta A, Aleman S, Cornillet M, Hertwig L, Strunz B, Lentini A, Reinius B, Brownlie D, et al. Natural killer cell immunotypes related to COVID- 19 disease severity. Sci Immunol. 2020;5(50).

30. Mooney JP, Qendro T, Keith M, Philbey AW, Groves HT, Tregoning JS, Goodier MR, and Riley EM. Natural Killer Cells Dampen the Pathogenic Features of Recall Responses to Influenza Infection. Front Immunol. 2020;11(135.

31. Chen G, Wu D, Guo W, Cao Y, Huang D, Wang H, Wang T, Zhang X, Chen H, Yu H, et al. Clinical and immunological features of severe and moderate coronavirus disease 2019. J Clin Invest. 2020;130(5):2620–9.

32. Tan L, Wang Q, Zhang D, Ding J, Huang Q, Tang YQ, Wang Q, and Miao H. Lymphopenia predicts disease severity of COVID-19: a descriptive and predictive study. Signal Transduct Target Ther. 2020;5(1):33.

33. Grifoni A, Weiskopf D, Ramirez SI, Mateus J, Dan JM, Moderbacher CR, Rawlings SA, Sutherland A, Premkumar L, Jadi RS, et al. Targets of T Cell Responses to SARS-CoV- 2 Coronavirus in Humans with COVID-19 Disease and Unexposed Individuals. Cell. 2020;181(7):1489–501 e15.

34. Braun J, Loyal L, Frentsch M, Wendisch D, Georg P, Kurth F, Hippenstiel S, Dingeldey M, Kruse B, Fauchere F, et al. SARS-CoV-2-reactive T cells in healthy donors and patients with COVID-19. Nature. 2020.

35. Lum JJ, DeBerardinis RJ, and Thompson CB. Autophagy in metazoans: cell survival in the land of plenty. Nat Rev Mol Cell Biol. 2005;6(6):439–48.

36. Pua HH, Guo J, Komatsu M, and He YW. Autophagy is essential for mitochondrial clearance in mature T lymphocytes. J Immunol. 2009;182(7):4046–55.

37. Xu X, Araki K, Li S, Han JH, Ye L, Tan WG, Konieczny BT, Bruinsma MW, Martinez J, Pearce EL, et al. Autophagy is essential for effector CD8(+) T cell survival and memory formation. Nat Immunol. 2014;15(12):1152–61.

38. Akbar AN, and Vukmanovic-Stejic M. Telomerase in T lymphocytes: use it and lose it? J Immunol. 2007;178(11):6689–94.

39. Mathew D, Giles JR, Baxter AE, Oldridge DA, Greenplate AR, Wu JE, Alanio C, Kuri- Cervantes L, Pampena MB, D’Andrea K, et al. Deep immune profiling of COVID-19 patients reveals distinct immunotypes with therapeutic implications. Science. 2020;369(6508).

40. Laing AG, Lorenc A, Del Molino Del Barrio I, Das A, Fish M, Monin L, Munoz-Ruiz M, McKenzie DR, Hayday TS, Francos-Quijorna I, et al. A dynamic COVID-19 immune signature includes associations with poor prognosis. Nat Med. 2020;26(10):1623–35.

41. Zhou R, To KK, Wong YC, Liu L, Zhou B, Li X, Huang H, Mo Y, Luk TY, Lau TT, et al. Acute SARS-CoV-2 Infection Impairs Dendritic Cell and T Cell Responses. Immunity. 2020;53(4):864–77 e5.

42. Cervantes-Barragan L, Zust R, Weber F, Spiegel M, Lang KS, Akira S, Thiel V, and Ludewig B. Control of coronavirus infection through plasmacytoid dendritic-cell-derived type I interferon. Blood. 2007;109(3):1131–7.

43. Langlois RA, and Legge KL. Plasmacytoid dendritic cells enhance mortality during lethal influenza infections by eliminating virus-specific CD8 T cells. J Immunol. 2010;184(8):4440–6.

44. Wolf AI, Buehler D, Hensley SE, Cavanagh LL, Wherry EJ, Kastner P, Chan S, and Weninger W. Plasmacytoid dendritic cells are dispensable during primary influenza virus infection. J Immunol. 2009;182(2):871–9.

45. Unterman A, Sumida TS, Nouri N, Yan X, Zhao AY, Gasque V, Schupp JC, Asashima H, Liu Y, Cosme C, et al. Single-Cell Omics Reveals Dyssynchrony of the Innate and Adaptive Immune System in Progressive COVID-19. medRxiv. 2020:2020.07.16.20153437.

46. Wilk AJ, Rustagi A, Zhao NQ, Roque J, Martinez-Colon GJ, McKechnie JL, Ivison GT, Ranganath T, Vergara R, Hollis T, et al. A single-cell atlas of the peripheral immune response in patients with severe COVID-19. Nat Med. 2020;26(7):1070–6.

47. McGill J, Van Rooijen N, and Legge KL. IL-15 trans-presentation by pulmonary dendritic cells promotes effector CD8 T cell survival during influenza virus infection. J Exp Med. 2010;207(3):521–34.

48. Wolfel R, Corman VM, Guggemos W, Seilmaier M, Zange S, Muller MA, Niemeyer D, Jones TC, Vollmar P, Rothe C, et al. Virological assessment of hospitalized patients with COVID-2019. Nature. 2020;581(7809):465–9.

49. Stuart T, Butler A, Hoffman P, Hafemeister C, Papalexi E, Mauck WM, 3rd, Hao Y, Stoeckius M, Smibert P, and Satija R. Comprehensive Integration of Single-Cell Data. Cell. 2019;177(7):1888–902 e21.

50. Becht E, McInnes L, Healy J, Dutertre CA, Kwok IWH, Ng LG, Ginhoux F, and Newell EW. Dimensionality reduction for visualizing single-cell data using UMAP. Nat Biotechnol. 2018.

51. Sun H, Zhou Y, Fei L, Chen H, and Guo G. scMCA: A Tool to Define Mouse Cell Types Based on Single-Cell Digital Expression. Methods Mol Biol. 2019;1935(91-6.

52. Zilionis R, Engblom C, Pfirschke C, Savova V, Zemmour D, Saatcioglu HD, Krishnan I, Maroni G, Meyerovitz CV, Kerwin CM, et al. Single-Cell Transcriptomics of Human and Mouse Lung Cancers Reveals Conserved Myeloid Populations across Individuals and Species. Immunity. 2019;50(5):1317–34 e10.

53. Greene CS, Krishnan A, Wong AK, Ricciotti E, Zelaya RA, Himmelstein DS, Zhang R, Hartmann BM, Zaslavsky E, Sealfon SC, et al. Understanding multicellular function and disease with human tissue-specific networks. Nat Genet. 2015;47(6):569–76.

